# Learning directed acyclic graphs for ligands and receptors based on spatially resolved transcriptomic analysis of ovarian cancer

**DOI:** 10.1101/2021.08.03.454931

**Authors:** Shrabanti Chowdhury, Sammy Ferri-Borgogno, Anna P Calinawan, Peng Yang, Wenyi Wang, Jie Peng, Samuel C Mok, Pei Wang

## Abstract

To unravel the mechanism of immune activation and suppression within tumors, a critical step is to identify transcriptional signals governing cell-cell communication between tumor and immune/stromal cells in the tumor microenvironment. Central to this communication are interactions between secreted ligands and cell-surface receptors, creating a highly connected signaling network among cells. Recent advancement in in situ-omics profiling, particularly spatial transcriptomic (ST) technology, provide unique opportunities to directly characterize ligand-receptor signaling networks that powers cell-cell communication. In this paper, we propose a novel statistical method, `LRnetST`, to characterize the ligand-receptor interaction networks between adjacent tumor and stroma cells based on ST data. `LRnetST` utilizes a directed acyclic graph (DAG) model with a novel treatment to handle the zero-inflated distribution observed in the ST data. It also leverages existing ligand-receptor regulation databases as prior information, and employs a bootstrap aggregation strategy to achieve robust network estimation. Application of `LRnetST` to ST data of high-grade serous ovarian tumor samples revealed both common and distinct ligand-receptor regulations across different tumors. Some of these interactions were validated through a MERFISH data set of independent ovarian tumor samples. These results cast light on biological processes relating to the communication between tumor and immune/stromal cells in ovarian tumors. An open-source R package of `LRnetST` is available on GitHub at https://github.com/jie108/LRnetST.

## Introduction

High grade serous ovarian cancer (HGSC) is the most lethal gynecologic malignancy, with its daunting overall survival rate showing limited improvement over decades [1, 2, 3, 4]. A major obstacle in fully understanding the mechanisms of tumor progression and chemo-resistance in HGSC is its high intra-tumor heterogeneity, comprising both tumor clonal heterogeneity and tissue architecture heterogeneity [5]. The latter is reflected by the heterogeneous stromal and immune cell population in the ovarian tumor micro-environment (TME) [6, 5]. The recent advances in in situ omics analysis have suggested an important link between cell-cell interactions among tumor/immune/stromal cells in TME and tumor progression as well as therapeutic resistance [7]. However, the molecular mechanisms that shape these cell-cell interactions in HGSC are still largely unexplored.

A predominant form of cell-cell signaling is powered through interactions between ligands from one cell and cognate receptors on neighboring cells. Considerable effort has been dedicated to develop tools for exploring these interactions using single-cell RNA-seq (scRNA-seq) data, including CellphoneDB [8], Single-CellSignalR [9], Cellchat [10], ICELLNET [11], CrosstalkeR [12], Nichenet [13], scMLnet [14], and CytoTalk [15]. Despite these endeavors to characterizing intercellular communications with scRNAseq data, the absence of spatial information in scRNAseq data significantly hampers the precision in dissecting ligand-receptor interaction network, given that these interactions occur locally between neighboring cells within the tissue.

The latest development of spatial transcriptomic (ST) profiling technology enables one to map mRNA molecules to a specific location (spot or grid) of a tissue slide in a high-throughput manner [16, 17, 18]. These platforms thus provide an unprecedented opportunity to comprehensively characterize the ligand-receptor interactions among neighboring cells (e.g. those from the adjacent grids on ST slides), which is not feasible based on either bulk or single-cell RNA profiles. In this paper, we aim to characterize the ligand-receptor regulatory network in HGSC using spatial transcriptomic data.

Pioneering effort have been made for inferring cell-cell communication networks while considering spatial information, including `Giotto` [19] and `stLearn` [20]. `Giotto`, a comprehensive open-source toolbox, is designed for analyzing and visualizing spatial transcriptomic data. Within `Giotto`, the Spatially Informed Cell-to-Cell Communication (*spatCellCellcom*) module calculates cell-cell communication scores for each ligand-receptor (L-R) pair between proximal cell types according to the spatial network. Permutation-based p-values and multiple hypotheses adjustment are subsequently computed for each ligand-receptor pair. On the other hand, `stLearn` focuses on the proportion of neighboring spots with upregulated expressions for both the ligand and receptor genes of a given L-R pair. The significant L-R pairs are then obtained by integrating the signals across all spots. Although, both methods incorporate spatial information in the inference of cell-cell communication, neither directly addresses the issue of zero-inflation that is pervasive in ST data. Furthermore, assessing co-expression based on either thresh-holding or marginal correlation is susceptible to the considerable variability present in ST data, leading to a lack of reproducibility, as demonstrated in numerical examples (see Results section).

To address these challenges, we developed `LRnetST` - <u>L</u>igand-<u>R</u>eceptor <u>Net</u>work learning based on <u>S</u>patial <u>T</u>ranscriptomics data, a novel tool to construct ligand-receptor networks between adjacent cells of different types based on either multi-cellular or single-cell ST data. The `LRnetST` pipeline begins by introducing the Neighbor Integrated Matrix (NIM), which integrates the spatial information and the molecular information within the ST data. Subsequently, `LRnetST` utilizes a binary variable along-side a continuous variable for every ligand/receptor in the node space. This coding strategy not only addresses zero inflation in ST data, but also enhances the power to detect interactions predominantly signaled through active/inactive statuses of ligands/receptors. To account for variation in library sizes across grids/cells in one ST experiment, `LRnetST` employs an aggregation framework that combines a bootstrap (bagging) procedure of DAG learning with down-sampling based normalization. This aggregation framework provides better control over false edge detection. Moreover, the DAG construction step in `LRnetST` is adaptable to incorporate prior information such as data from existing ligand-receptor databases.

We applied LRnetST to both multi-cellular and single-cellular ST datasets of ovarian cancer. In the multi-cellular dataset, our focus was on detecting ligand-receptor interactions between neighboring spot-pairs enriched with tumor and stroma cells, respectively. In the single-cellular ST data, we examined interactions between tumor cells and other immune or stroma cell types. Specifically, we first applied the proposed LRnetST pipeline to a 10X Genomics ST data of 4 HGSC samples. To guide the construction of DAGs, we incorporate known ligand-receptor regulation information from relevant databases [22] as prior information to constrain the network space. The LRnetST analysis revealed a substantial number of shared ligand-receptor interactions between tumor and stroma cells across four different HGSC samples, outperforming alternative methods. Notably, both LRP1 and ITGB1 were identified as hub nodes, showing connections to multiple ligands in adjacent regions within each of the four patient-specific LR networks. Further applying LRnetST to an independent single-cell MERFISH ST data of ovarian tumors, we were able to confirm several LR interactions, including those between LRP1 in tumor cells and the protease SERPINA1 in the neighboring stromal cells, as well as interactions involving ITGB1 in tumor cells with VEGFA and VCAM1 in neighboring stromal cells. These findings offer fresh insights into the roles of these tumor-relevant genes/proteins in facilitating cell-cell communication within HGSC.

## Materials and methods

### Notation and background

We first provide a brief review of directed acyclic graph (DAG) models. A directed acyclic graph 𝒢 (ℕ, 𝔼) contains a node set ℕ and an edge set 𝔼, where the edges are directed from *parent* nodes to *children* nodes, without any cycle in the graph. In the DAG models, the node set ℕ represents a set of random variables, while the edge set 𝔼 represents the conditional dependence relationships among these random variables. Structure learning of a DAG means identifying the parent set (often known as neighborhood) of each node in the graph. Different DAGs could represent the same set of conditional dependencies, i.e., they form an equivalent class of DAGs. It is shown that, two DAGs are equivalent if and only if they have the same set of skeleton edges (obtained by removing directions from the directed edges) and *v*-structures ([21]). A *v*-structure is a triplet of nodes (*w*_1_, *w*_3_, *w*_2_), such that *w*_1_ → *w*_3_ ∈ 𝔼, *w*_2_ → *w*_3_ ∈ 𝔼, and *w*_1_, *w*_2_ are not adjacent. Primarily, there are three classes of methods for DAG structure learning: (1) *score-based* methods (e.g., [22]), (2) *constraint-based* methods, e.g., PC algorithm (`PC-alg`) [21, 23, 24] and (3) *hybrid* methods, e.g., Max-Min Hill Climbing (`MMHC`) [25].

### LRnetST

We present a new tool - `LRnetST` - for learning ligand-receptor interaction networks based on spatially resolved transcriptomic data. First, `LRnetST` integrates the spatial and the molecular information in the ST data by introducing Neighbor Integrated Matrices (NIM). Second, to account for the high dropout rates in the ST data, `LRnetST` codes each gene’s active/inactive (expressed/not-expressed) status by a binary variable and uses a continuous variable to represent one gene’s expression level given the gene is active (expressed). Moreover, to account for the varying UMI count levels across different ST spots, `LRnetST` incorporates an aggregation framework that conveniently couples the downsampling based normalization with a bootstrap based network inference procedure.

Finally, `LRnetST` can take into account prior information of interactions, such as those from existing ligand-receptor databases by constraining the DAG search space to the documented edges among the ligands and receptors in the databases. **Figure** 1 illustrates the flow of the `LRnetST` pipeline. See also **Algorithm** in section A.1 in the Supplementary Material. In the following, we elaborate the major steps.

#### Neighbor Integrated Matrix (NIM)

In the `LRnetST` pipeline, we first derive an initial Neighbour Integrated Matrix (Initial-NIM) to integrate the spatial and the molecular information in ST data. Since our goal is to characterize whether/how receptors in non-tumor cells in the micro-environment are affected by the ligands from the neighboring tumor cells or vice versa, we choose to focus on pairs of adjacent ST spots enriched of tumor and stromal/immune cells, respectively. Specifically, for each tumor sample, ST spots are first classified into two classes: enriched or not enriched of tumor cells. We then identify the subset of tumor-cell-enriched spots sitting on the boundary of the tumor regions (i.e. being adjacent to spots enriched of non-tumor cells on the ST slice). We refer to these spots as the Index Spots and refer to their closest non-tumor-cell-enriched spots as their Neighbor Spots. Denote the total number of Index Spot and Neighbor Spot pairs as *n*, and the number of genes under consideration as *p*.

In the next step, we derive an Initial-NIM by stacking the rows of the gene expression matrix of the Index Spots, which has a dimension of *p×n*, with the gene expression matrix of the corresponding Neighbor Spots, which also has a dimension of *p× n*. The resulting Initial-NIM has an expanded feature space of 2*p* rows (**Figure** 1). Finally, we convert the *Initial-NIM* into *NIM* by replacing the expression *Z* of a gene with a pair of features, *X* and *Y* to facilitate the modeling of zero-inflation in the ST data (Please see the next subsection on “Accounting for zero inflation in ST data”). The resulting NIM has a dimension 4*p × n*. See **Figure 1B** and the pseudo code of the algorithm outlined in section **A.1** of the Supplementary Material for more details.

**Figure 1.**
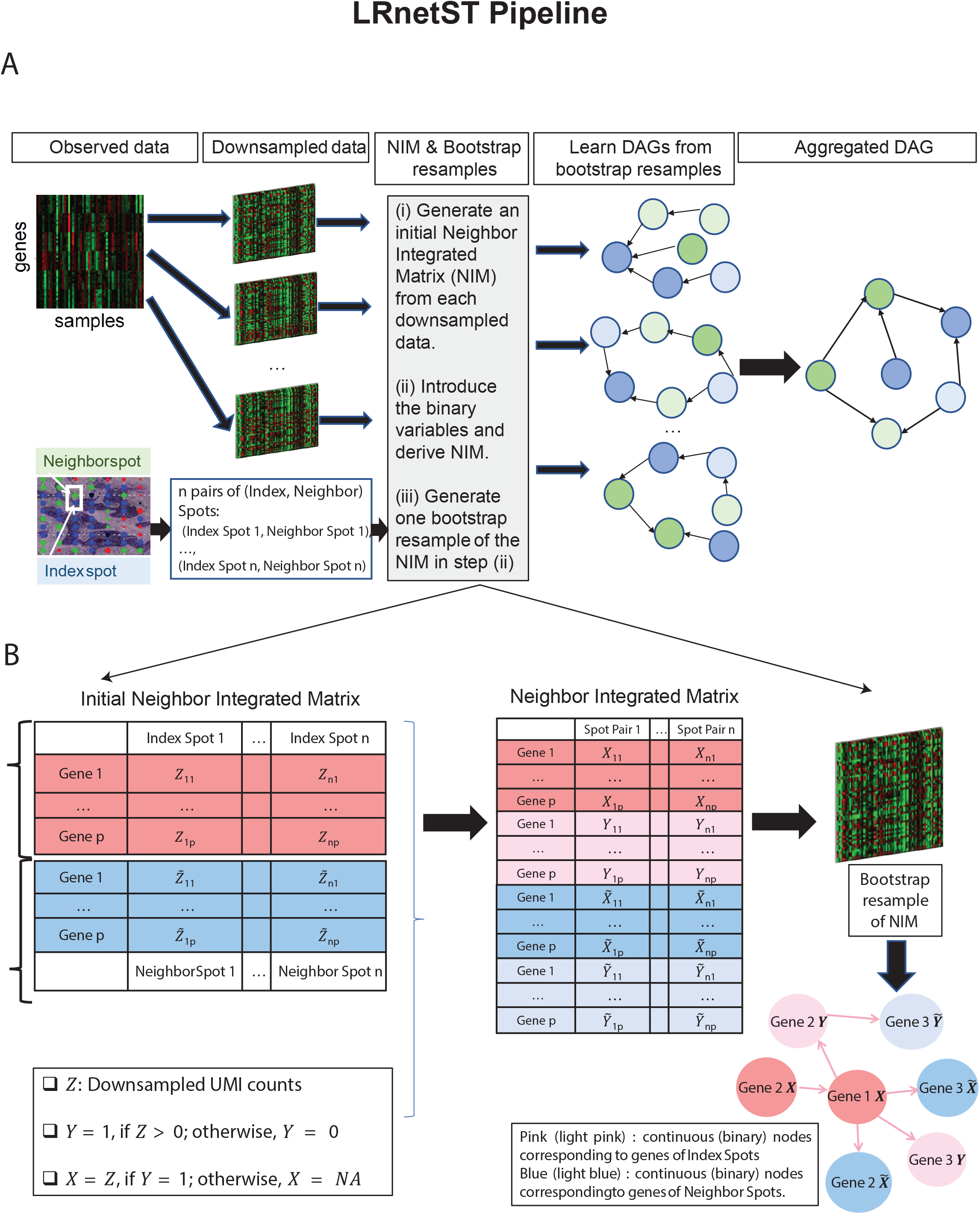

#### Accounting for zero inflation in ST data

Similar to scRNA-seq data, ST data has a high dropout rate, i.e. only a fraction of the transcriptome can be detected in the sequencing result of each ST spot. To facilitate the modeling of such zero-inflated distributions, `LRnetST` uses two nodes (variables) to represent each gene: a binary node and a continuous node.

First, we use *Z*_*ij*_ to denote the downsampled UMI count of the *i*_*th*_ gene in the *j*_*th*_ ST spot (see *Section 3.2.4* for details on downsampling normalization). Note, *Z*_*ij*_ = 0 usually corresponds to the situation where either the *i*_*th*_ gene is not expressed, or its expression level is low in the *j*_*th*_ spot and thus is difficult to be detected in the sequencing experiments. Therefore, we view the event of *Z*_*ij*_ = 0 as a “less-active” status of the gene and model this status directly. Specifically, the binary node, denoted as *Y*_*ij*_, represents the detection status of a gene:

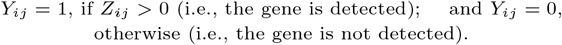

Given the gene is detected. (i.e., *Y*_*ij*_ = 1), the continuous node, denoted as *X*_*ij*_, represents the normalized gene expression/abundance:

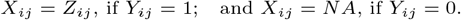

Then, by including both nodes, *X*_*ij*_ and *Y*_*ij*_, `LRnetST` not only nicely accounts for the zero inflation in the data distribution, but also achieves better power in detecting those interactions that are largely signaled through the active/less-active statuses of genes.

#### Downsampling normalization and bootstrap model aggregation

`LRnetST` also employs an aggregating framework that couples a bootstrap (bagging) procedure of DAG learning with the downsampling based normalization to achieve better control of false edge detection and to account for the varying library sizes or cell type compositions across different tissue spots.

We apply downsampling normalization on the original gene expression data by fixing the total UMI at the median level of the total UMI counts across all ST spots. We then generate *B* = 100 downsampled gene expression matrices ([*Z*]), and construct *B* = 100 NIMs; also see **Figure 1(b)**) accordingly. We then obtain one bootstrap resample on each NIM through sampling with replacement of the Index/Neighbor Spots pairs (i.e., the columns of the NIM). Finally, we learn one DAG based on each bootstrap resample using the method described in the previous subsection.

The resulting ensemble of DAGs are then aggregated following the aggregation procedure implemented in `DAGBagM` [26], where the hill climbing algorithm is (again) used to search for the DAG that minimizes an aggregation score based on the *structural Hamming distance (SHD)* while maintaining acyclicity.

#### Score function and optimization

In `LRnetST`, we learn DAG by minimizing a score function through a greedy search algorithm (*hill-climbing*), where the score of the continuous nodes are calculated only using data from spots on which their binary nodes take the value 1 (i.e., *X* is not *NA*). This makes the binary nodes natural parents of the corresponding continuous nodes of the same gene. At each search step, the score of a continuous node, *X*, is calculated by regressing *X* onto its current parent set, using only the samples where *X* is non-zero: For a given graph *𝒢*, the BIC score is calculated as 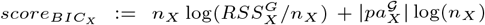, where, 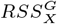 is the residual sum of squares, *n*_*X*_ is the number of samples where *X* is non-zero and 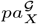 denotes the parent set of *X* in graph 𝒢. The score of a binary node *Y*, is obtained by regressing *Y* onto its current parent(s) 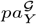 through logistic regression:

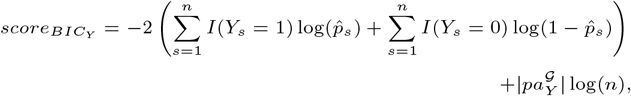

where 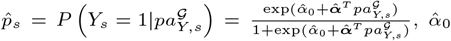 is the fitted intercept and 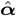 is the vector of the fitted coefficients in logistic regression, and *n* is the sample size.

The final score of a graph 𝒢 is the summation of the scores of the individual nodes:

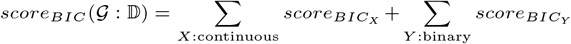

where, 𝔻 denotes the data. We then search for the DAG that minimizes the above score function using an efficient implementation of the hill climbing algorithm that is modified from one of our recent works `DAGBagM` [26] – a DAG method for mixed types of nodes.

### Simulation Studies

We utilized synthetic data sets to evaluate the performance of the DAG construction step in `LRnetST`. When generating the synthetic data sets, we considered various scenarios with different numbers of genes, (true) edges, and Index-Neighbor spot pairs (i.e., samples sizes) (see below). We compared the performance of DAG construction based on initial-NIM and NIM to assess the advantage of using the paired binary and continuous nodes to facilitate the modeling of zero-inflation in ST data. Specifically, we compared the performance of `LRnetST` with three alternative methods: `DAGBagM`, `DAGBagM C` [26] and `bnlearn` [27]. `DAGBagM C` and `bnlearn` were applied to the Initial Neighbor Integrated Matrix (**Figure** 1b) before introducing the binary nodes. Note that `bnlearn` was applied to initial-NIM after converting it to binary variables using median cut-off. `DAGbagM`, similar to `LRnetST`, is applied to the NIM after introducing the binary variables in the node space. However, for `DAGbagM`, *X* is replaced with *Z* (*NA* in *X* is replaced with 0) in NIM, and all samples are used to calculate the scores for the continuous nodes; whereas `LRnetST` uses the converted *X* so that only samples with *Y* = 1 are used to calculate the scores for the continuous nodes. Note that, we performed bootstrap for all methods except bnlearn due to its prohibitive computational cost.

We consider simulations with different sample sizes (i.e. numbers of Index-Neighbor Spot pairs: *n* = 200 and 400) and varied the number of genes (*p* = 100, 200, 300, 400, 500, and 600) with different number of true edge, thus generating 12 DAG topology. The true DAGs corresponding to different *p* are shown in **Figure** S1A-F.

Given *p* genes, we first introduce 4*p* nodes as following: For a given gene *i*, we use two nodes (*Z*_*i*_, *Y*_*i*_) to indicate its expression level and active status in the Index Spot, and 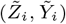 to indicate its expression level and active status in the Neighbor Spot. We then randomly choose *m* edges (while excluding cycles) between one node corresponding to Index Spot and another node corresponding to Neighbor Spot. The edges are of the following patterns: (i) 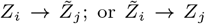 or (ii) 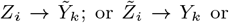 (iii) 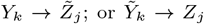or (iv) 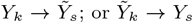. This process creates a DAG 𝒢^*^ with *p*^*^ = 4*p* nodes and *m* edges. We also introduce directed edges from each binary node 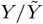 to its corresponding continuous node 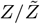 and pass them as *whitelist* (i.e., edges always kept in the search steps) while learning the DAGs using `LRnetST` and `LRnetST.naive`.

Given 𝒢^*^ and sample size *n*, i.i.d. samples of continuous nodes are generated according to *Gaussian linear mechanisms*: 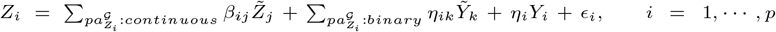, where *ϵ*_*i*_’s are independent Gaussian random variables with mean zero and variance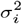. The coefficients *β*_*ij*_ s, *η*_*ij*_ s, etc., are uniformly generated from ℬ = [−0.5, −0.3] ∪ [0.3, 0.5] and the noise variances 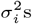 are chosen such that for each node the corresponding *signal-to-noise-ratio (SNR)*, defined as the ratio between the standard deviation of the signal part and that of the noise part, is within [0.5, 1.5]. Moreover, i.i.d. samples for the binary nodes are generated by logistic regression models: 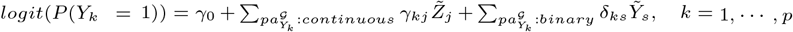. Binary nodes without any parent are generated from Bernoulli(0.5). 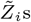 and 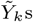 are generated similarly. Finally, for each gene 1 ≤ *i* ≤ *p* and on each sample 1 ≤ *l* ≤ *n*, we define *X*_*il*_ = *Z*_*il*_ if *Y*_*il*_ = 1; Otherwise 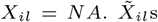 are defined similarly.

For a given sample size, we generate 100 independent data sets for each DAG. For each simulated dataset, we fit `LRnetST` to the 4*p × n* stacked *X*, 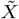, *Y* and 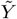 matrix (see **Figure** 1) and obtain the estimated DAG. For the purpose of comparison, `DAGBagM` is applied to the 4*p × n* stacked *Z*, 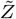, *Y* and 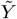 matrix, with the same *whitelist* used as mentioned above. On the other hand, `DAGBagM` and `bnlearn` are applied to the 2*p × n* stacked *Z* and 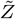 matrix with only continuous nodes. Note that, `DAGBagM` is a previously developed algorithm by us where the same aggregation procedure has been implemented as in `LRnetST`; whereas we use `bnlearn` without any aggregation (due to its computational slowness). Both `DAGBagM` and `bnlearn` are score-based DAG learning methods using the hill-climbing algorithm.

We then assessed the powers (or true positive rates), false discovery rates (FDRs) and the F1 scores to evaluate the performance of `LRnetST` and other methods for (i) detecting the skeleton edges (i.e., edges without direction); and (ii) detecting the directed edges. Power (or TPR) is calculated as the ratio of the number of edges identified correctly in the estimated DAG and the number of total edges in the true DAG. FDR is calculated as the ratio of the number of edges that are falsely identified in the estimated DAG and the number of total edges in the estimated DAG. F1-score is calculated as *F* 1 = 2 *×* precision *×* recall*/*(precision+recall), where precision=1-FDR and recall = power. The performance of each method under different scenarios are presented in Table A.1 in Supplementary Material, where all numbers are averaged over the 100 independent replicates.

### Data Description and Processing

#### 10X Genomics ST Data

Spatial transcriptomic analysis was performed on four fresh frozen treatment-naive advanced stage HGSC samples using the 10X Genomics platform as described in our previous work [28]. Among these samples, two (A4 and A5) were derived from chemo-sensitive patients with extended progression free survival time (¿10 years); and the remaining two (A10 and A12) originated from chemo-resistant patients with short progression free survival time (¡6 months). In the ST experiment, each ST spot contained 20-50 cells, and 930-1007 spots were profiled per tumor slice.

Expression levels of genes in each spot were measured using counts of unique molecular identifier (UMI). Specifically, the output UMI count matrix for each sample (spot) is obtained using ST pipeline (version 1.80.1) following the recommended default parameter setting from FASTQ files [29]. Human genome reference (GRCh38) was used for read alignment and corresponding genome index was generated using STAR2 (version 2.7.9a) [30]. In the preprocessed ST data, the number of genes with non-zero UMI ranged from 68 - 9364 across all grids in all four samples. The median of the total-UMI per grid is 87633.5, 1066, 13318, and 7473 for the four tumor samples, respectively.

For the 10X genomics ST datasets, since each spot is a mixture of several cell types, in order to identify tumor or immune/stromal cell-enriched spots, we estimated the cell-type proportions in each grid (spot) by performing deconvolution analysis using the function *runDWLSDeconv* implemented under `Giotto` with the scRNAesq data of ovarian tumors from from five HGSC patients [31] as the reference. Specifically, we used the gene signatures upregulated in the cell-types Adipose, BCL, Endothelial, Epithelial, Fibroblast, Macrophage, Monocyte, NKC and TC8 from the above-mentioned scRNAseq data and used the *R* function *runDWLSDeconv* implemented under `Giotto` to estimate the cell-type proportions in the 10X ST data. Based on the proportions, we only kept the spots with (1) high Epithelial cell percentage (low immune/stromal cell percentage) and (2) high immune/stromal cell percentage (low Epithelial cell percentage) and filtered out the remaining. Further we inspected the H&E image of the tissues to annotate the tumor enriched spots from (1) and spots enriched with immune/stromal cells from (2).

For each tumor spot, we further identified the adjacent or neighboring immune/stromal spot that are at most 2-grid distance apart from the tumor spot. In this way, we filtered out some of the tumor spots that did not have any immune/stromal spot within 2-grid distance.

We only considered the genes that were documented in the ligand-receptor interaction database, and further filtered out those detected in less than 20 % of the spots in one tumor tissue.

#### MERFISH ST Data

MERFISH data sets for four tumor samples (one tumor slice from one patient and three tumor slices from another patient) were downloaded from <u>this website</u> [32]. Unlike the 10X Genomics platform, MERFISH is a spatially resolved single-cell transcriptome profiling technology. For each tumor slice, before normalization, we filtered out the cells with less than 10 genes observed and filtered out the genes that were observed in less than 1500 cells. We then performed library-size normalization using the function *normalizeGiotto* implemented in `Giotto R` package. The average number of genes per cell ranged from 10 to 346 in the four tumor slices, while the number of cells after filtering were 355633, 248065, 68953 and 209173 in the four tumor slices, respectively.

After pre-processing using the pipeline implemented under `Giotto` [19] including normalization and filtering of the cells with less then 10 genes measured and filtering of genes measured in less than 1500 cells, we clustered the cells of each tumor tissue using *doLeidenCluster* implemented under `Giotto`. We used the same single cell data as used for deconvolution of the spots in 10X ST data [31] to identify the marker genes upregulated in each of the 9 cell types mentioned above. We mapped the marker genes to the clusters in the MERFISH data, e.g., we combined the cells from all the clusters where the Epithelial marker genes were upregulated as defined the new class as tumor cluster, and so on. This way we identified the groups of cells as tumor, macrophage, T-cells, fibroblast or endothelial in each of the four tumor tissues (Table S2). The processed data with cell-type labeling was then used for LR network construction and validation.

To identify the cells from different cell types that are closest to the tumor cells, we constructed a Delaunay network using the function *createSpatialNetwork* implemented under `Giotto`. For each tumor cell, we identified the cell adjacent/neighboring to the tumor cell that has minimum distance from the tumor cell, obtained from the network.

As in the 10X genomics ST data, for the network learning in MERFISH data also, we only considered the genes that were documented in the ligand-receptor interaction database, and further filtered out those detected in less than 10 % of the cells in one tumor tissue. The median of the total UMIs per cell in each sample was used for the downsampling normalization in `LRnetST`. Table A.3 in the Supplementary Material gives the detailed numbers of the tumor and neighboring cells, as well as the numbers of adjacent index-neighbor-cell pairs in each tumor tissue.

## Results

### LRnetST pipeline

To effectively characterize the complicated ligand-receptor regulatory relationship in TME, we proposed to use directed acyclic graphs (DAGs), which has emerged as a valuable tool for inferring gene-gene interactions [33, 34, 35, 36, 37, 38, 39]. However, multiple challenges impede the application of existing DAG models to ST data for constructing ligand-receptor networks. Firstly, integrating both spatial information and -omics profiles concurrently into ligand-receptor network learning is not readily addressed by current DAG methods/software. Secondly, in the sequencing output of an ST experiment, a substantial portion of transcripts is often missing due to a combination of both biological factors (e.g., absence of gene expression in the cell population of that grid) and technical limitations (e.g. limited read depth). The resulting zero-inflated distributions of Unique Molecular Identifier (UMI) counts impose statistical and computational challenges in network inference and mechanistic interpretation. Thirdly, UMI count levels of different spots on one tissue slice are subject to both technical variations such as different library sizes, and biological variations such as cell type heterogeneity. Consequently, proper normalization across spots is essential before undertaking downstream analysis.

To address these challenges, we developed a new DAG model coupled with customized fitting pipeline, called `LRnetST`, for constructing ligand-receptor networks based on ST data. Here, we provide a concise overview of the key steps in the `LRnetST` pipeline, with comprehensive details of the statistical models available in the Methods section and the complete algorithm pseudocode in Section A.1 of the Supplementary Material.

As illustrated in Figure 1A, the `LRnetST` pipeline comprises of four major steps: (1) normalization through down-sampling, (2) construction of the Neighbor Integrated Matrices (NIM) with an additional round of sample bootstrap, (3) estimation of DAGs based on bootstrapped samples, and (4) aggregation of the DAGs into a final network.

Within the `LRnetST` pipeline, the second step contains three key components (Figure 1B). Firstly, for a given normalized ST data matrix (gene by spot/cell) obtained through down-sampling, `LRnetST` integrates the spatial and molecular information through introducing the Initial Neighbor Integrated Matrices (Initial-NIM). This involves combining gene expression information from two neighboring spots/cells into one vector (column). Consequently, evaluating dependence between genes in neighboring spots/cells is equivalent to assessing the dependence between features from the top and bottom halves of the Initial-NIM matrices. Secondly, to account for the high dropout rates in the ST data, `LRnetST` encodes each gene expression using two variables: one binary indicator denoting the expressed or not-expressed status; and a continuous number representing the gene expression level when the UMI count is greater than 0. The resulting data matrix is referred to as NIM, whose feature space is doubled compared to that of the Initial-NIM. This strategy improves the power of detecting dependencies among genes, particularly in the presence of zero-inflated distributions, as demonstrated in the numerical examples below. The third component involves perturbing the sample space of the NIM through bootstrap, where the resulting matrix serves as the input of the subsequent DAG construction. Notably, the ensemble and aggregation framework in `LRnetST` through integrating down-sampling-based normalization, NIM construction, and the bootstrap-based network inference, not only effectively accounts for the varying UMI count levels across different ST spots/cells, but also enhances the robustness of the DAG estimation.

The third step of `LRnetST` contains a novel DAG model customized to the NIM matrices. Specifically, the DAGs are constructed by minimizing a score function, amounting to maximize a BIC penalized joint likelihood of all the random variables in the graph. Similar to a previous work [26], continuous variables conditioned on their parent nodes are modeled with Gaussian distributions, while a logistic model is employed to characterize the dependency of binary variables on their parent nodes. In addition, from the definition of the binary and continuous variable of each gene, it’s obvious that the binary variable is a parent node of its corresponding continuous variable and the likelihoods of the continuous variables are evaluated only for spots where the corresponding binary nodes take the value 1 (i.e., when the continuous variables are defined). Moreover, when building the DAG based on the NIM matrix, since edges of interests are between genes from different spots, only edges between features from the top (i.e., index spots) and the bottom (i.e., the paired neighbor spots) of the NIM matrix are considered. Moreover, `LRnetST` incorporates existing ligand-receptor databases as prior information to further constrain the search space of directed edges during DAG fitting by the hill climbing algorithm.

In the last step, the derivation of the final aggregated DAG is achieved by searching for a DAG that minimizes an average structural Hamming distance to the DAGs in the ensemble built from the previous steps (see Methods).

Please note that LRnetST is a versatile tool and can be applied to both multi-cellular and single-cellular ST datasets. In the real application detailed in this manuscript, we demonstrated LRnetST’s utility by analyzing both a multi-cellular and a single-cell ST datasets of ovarian cancer. For the multi-cellular data, our primary objective was to identify ligand-receptor interactions between neighboring spot-pairs enriched with tumor and stroma cells respectively. And to validate the findings, we extended the analysis to an independent single-cell ST dataset of ovarian cancer, where we investigated LR interactions between tumor cells and other immune or stroma cell types. For more details, please refer to the following sections.

### Evaluation of DAG construction based on synthetic data sets

We performed simulation studies to evaluate and compare the performance of the DAG construction step in `LRnetST` to alternatives including `DAGBagM` [26], `DAGBagM C` [26], and `bnlearn` (see the detail in Methods). The results in Table A.1 in Supplementary Material and Figure S1G-H indicate superior performance of `LRnetST` compared to the other methods across all considered scenarios, illustrating the advantages of including the binary nodes in NIM in the `LRnetST` pipeline. Furthermore, the application of both `LRnetST` and `DAGBagM` on NIM underscores the superior performance of `LRnetST` over `DAGBagM`, emphasizing the advantages derived from the customized treatment of likelihood terms associated with paired binary and continuous nodes in `LRnetST`. In the end, as expected, for all four methods, we observed improved performances as the sample sizes (n) increased; while declined performance with the increase of the number of genes (p).

### Revealing cell-cell interaction in ovarian cancer with LRnetST

#### Applying LRnetST to the 10X Genomics ovarian cancer ST data

For each tumor sample in the 10X Genomic ovarian cancer ST data set (see the Data Description under Methods Section), we applied `LRnetST` to construct tumor-specific ligand-receptor networks. Firstly, to identify tumor or immune/stromal cell-enriched spots, we estimated the cell-type proportions in each grid (spot) by performing deconvolution analysis using *runDWLSDeconv* implemented in `Giotto` with the scRNAesq data of ovarian tumors from [31] as the reference (see Methods). Based on the cell-type proportion estimates, we then inspected the tissue and cell morphology in high resolution Hematoxylin and eosin stain (H&E) images, and manually annotated the ST grids (spots) enriched with tumor or stromal cells (Figure S2A-D, Methods).

To construct the NIM, we screened for tumor cell-enriched (index) spots with neighboring (within 2-grid-units on the ST slices) stromal cell-enriched spots. Using these index-tumor and neighboring-stroma spot-pairs, we assembled the Initial NIM and then NIM as described in the previous section (see Figure 1). Note, in this process, we focused on the genes documented in the ligand-receptor interaction database [40], and further filtered out those detected in less than 50% of the spots in one sample. The median of the total UMIs per spot in each sample was used for the down sampling normalization in `LRnetST`. Table A.2 in Supplementary Material gives the detailed numbers of the tumor and stromal spots, as well as the numbers of adjacent tumor-stromal-spot pairs in each sample. The table also lists the numbers of ligand/receptor genes with adequate UMI data points to construct the network in each sample.

Using `LRnetST`, we built four ligand-receptor networks and obtained 422, 170, 334, and 210 ligand-receptor regulation edges for samples A4, A5, A10 and A12, respectively (Figures S4A-B, S5A-B, Table S1). Both common and distinct interaction patterns were observed across the four tumor samples.

##### Comparing LRnetST with stLearn and spatCellCellcom

To compare the performance of `LRnetST` with other ligand-receptor network learning methods, we also applied `stLearn` and `spatCellCellcom` on the same data set. For pre-processing including normalization, we followed the respective pipelines of these two methods implemented in the corresponding R (`spatCellCellcom` implemented under `Giotto`) and python (`stLearn`) packages. Similar to `LRnetST`, we only considered the genes documented in the ligand-receptor database and survived the missing filtering cut-off of 50%. Both `stLearn` and `spatCellCellcom` provided adjusted p-values for all possible interactions between ligands and receptors. By thresh-holding the adjusted p-values, `stLearn` inferred 113, 111, 103 and 36 ligand-receptor regulation edges (Table S1), respectively; while `spatCellCellcom` identified 188, 123, 330 and 66 edges (Table S1), respectively, for samples A4, A5, A10 and A12.

##### Reproducibility of the edges across 4 tumor samples

We first evaluated how “reproducible” the inferred ligand-receptor (LR) edges are across different tumor samples. Intuitively, methods detecting more shared (reproducible) edges across tumors shall enjoy better power for detecting meaningful biological interactions. As summarized in Figure 2A-C (vertical bar heights indicating number of common edges shared across at least 2 out of 4 tumor samples), `LRnetST` inferred many more shared edges across multiple samples than the other two methods. Specifically, `LRnetST` inferred 185 common edges, while `spatCellCellcom` inferred 67 and `stLearn` inferred 48 common edges across the tumors. Note that, for counting the common edges, we considered undirected networks and further collapsed the binary and continuous component of the genes. For example, `LRnetST` identified 4 edges shared by all four tumors, whereas the other two methods failed to identify any such edges. Furthermore, we assessed whether the overlap between LR edges for a pair of tumors significantly surpassed what would be expected by random chance, as outlined in the Methods section. The resulting adjusted p-values are presented in Figure 2D. The outcomes from `LRnetST` exhibited significant overlap for all six pairs of tumors, whereas the results from `spatCellCellcom` and `stLearn` demonstrated significant overlap for only half of the pairs. This suggests the higher level of reproducibility of `LRnetST` compared to the alternatives.

**Figure 2.**
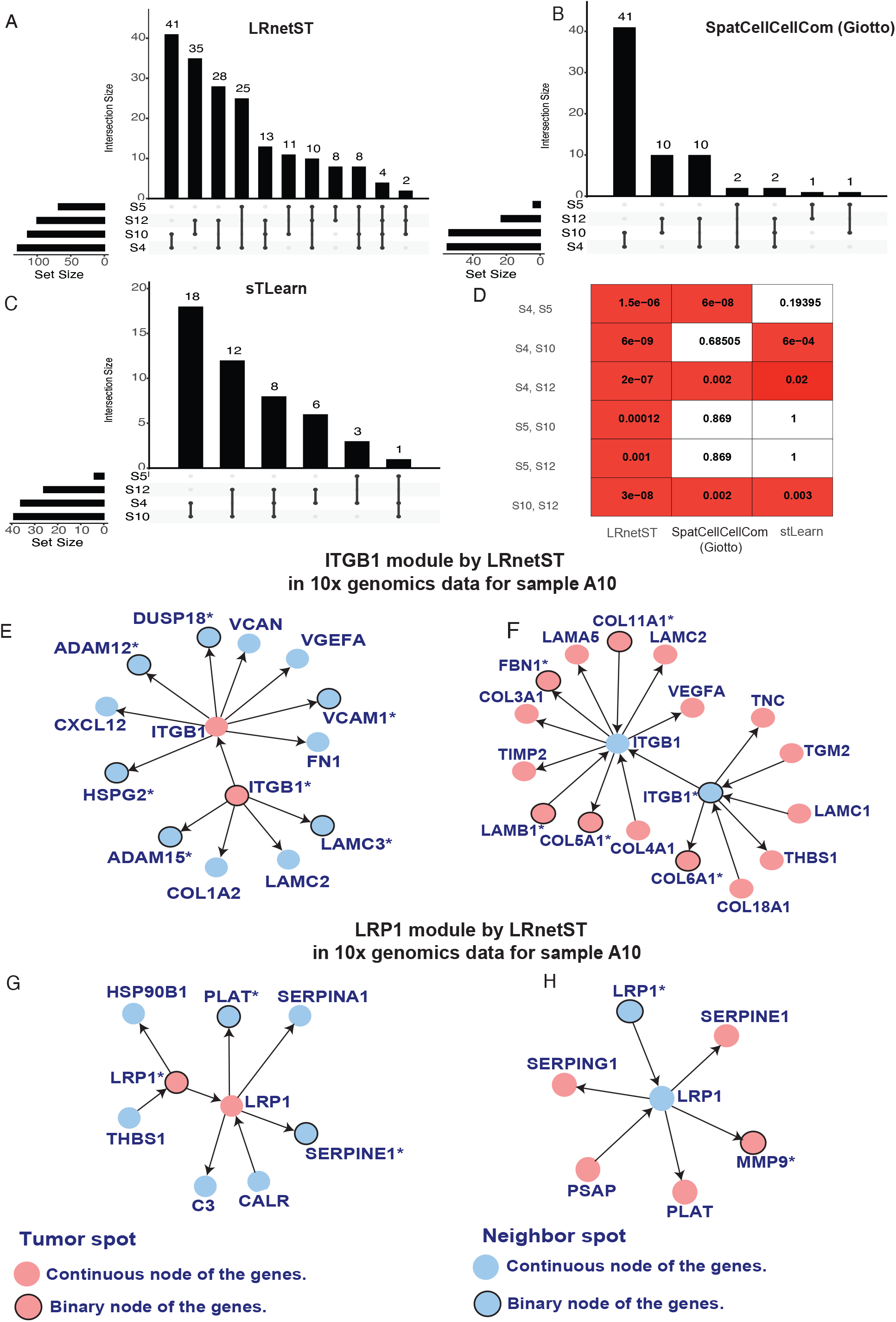

We then investigated the hub nodes in the LR networks by the three methods (Figure S3). Hub nodes are those with higher connections in a network, and thus are often more likely to play an essential role in the regulatory network [41]. Labeling the top 30 nodes with the largest degrees in each network as the hub nodes, we identified three common hub nodes, LRP1, ITGB1 and ITGAV, across all four tumors based on `LRnetST`’s results. On the contrary, only ITGB1 was detected as a common hub node by `spatCellCellcom` and `stLearn`.

Interestingly, in `LRnetST` results (Figure S3A), both LRP1 levels in tumor cell-enriched spots (LRP1 tumor) and that in the stroma cell-enriched spots (LRP1 stroma) appeared to be hub nodes in all LR networks. The degrees of the LRP1 tumor nodes are 8, 18, 7, and 4 in the four samples, respectively, while the degrees of the LRP1 stroma nodes are 13, 22, 5, and 4, respectively. Of all the edges associated with LRP1, the interaction between LRP1 stroma and MMP9 tumor (MMP9 levels in tumor cell-enriched spots) is the only common edge identified in all four tumor samples (Figure 2H and Figures S6B, D, F). Previous studies have highlighted the active involvement of the LRP1-MMP9 interaction in processes such as the migration of epithelial ovarian cancer (EOC) [42] and other cancer types, as well as in the development of cerebral edema (see Discussion).

Moreover, a few interactions involving LRP1 were exclusively detected in tumors from chemo-sensitive or chemo-refractory patients (Figure 2G-H, and Figure S6A-F). For example, in the two chemo-sensitive patients, LRP1 Tumor is inferred to interact with C1QB from the nearby stroma spots, which supports the potential role of LRP1-C1QB crosstalk network in modulating immune response in HGSC (see Discussion). On the other hand, LRP1 Tumor is detected to interact with SERPINA1, a serine protease inhibitor, from the nearby stroma spots only in samples from the two chemo-refractory patients. This regulation implies a potential connection between LRP1 and the protease [43].

Apart from LRP1, ITGB1 is another hub node inferred to interact with multiple genes in all four tumors (Figure 2E-F, and Figure S7A-D). Specifically, for both chemo-refractory patients, ITGB1 tumor was detected to interact with VCAM1 and VEGFA from the nearby stroma spots. At the same time, ITGB1 stroma was connected with multiple genes, including LAMC2 and VEGFA, from the nearby tumor spots. Multiple literatures reported that the intercellular communication happens through the interaction between ITGB1 and VEGFA [44]. In addition, ITGB1 and VCAM1 are both cell-adhesion molecules that influence immune response and anti-immunity [45] (see Discussion).

To further assess the performance of various methods, we then applied `LRnetST` to a cell-level MERFISH data to validate some of the detected LR interactions.

##### LRnetST applied on MERFISH data for validation

We studied four MERFISH data sets [32] derived from four tumor slices of two ovarian cancer patients (one slice from one patient and three slices from another patient). MERFISH, a spatially resolved single-cell transcriptome profiling technology, offers cell-level resolution, enabling detailed monitoring of LR interactions between individual cells. This granularity is particularly advantageous for overcoming the challenge of cell type mixtures within spots encountered in 10X genomics ST data. However, owing to sequencing limitations inherent to MERFISH, only a limited set of transcripts, in the range of few hundreds, can be detected and quantified. This restriction significantly constrains the ability to systematically characterize LR interactions, underscoring our use of MERFISH data for validation rather than exploration purposes.

For each tumor slice, after pre-processing, clustering and cell type labelling (see Methods), we identified neighboring tumor-stroma or tumor-immune cell pairs and constructed the NIM for applying `LRnetST`. The number of adjacent tumor-stromal/immune cell pairs for each tumor slice are given in Table A.3 in the Supplementary Material. Figure 3A illustrates the spatial distribution of tumor cells and macrophage cells in one tumor slice. While there are thousands of epithelial (pink) and macrophage (light blue) cells in the tumor slice, we focused on the neighboring tumor-macrophage pairs that are in proximity with each other (red epithelial cells and dark blue macrophage cells) for the `LRnetST` analysis. Similar plots illustrating spatial distribution of tumor cells and other stroma cell types, including T-cells, fibroblast and endothelial, are shown in Figure S8A-C. With `LRnetST`, we built a collection of ligand-receptor networks one for each cell type pair (i.e., tumor-macrophage, tumor-T-cell, tumor-fibroblast, and tumor-endothelial), and each tumor slice (Table S3). There were 53 nodes involved in LR interactions that were inferred in at least 2 tumor slices. Figure 3B-E illustrates subsets of the inferred networks considering different neighboring cell type pairs for one tumor slice.

**Figure 3.**
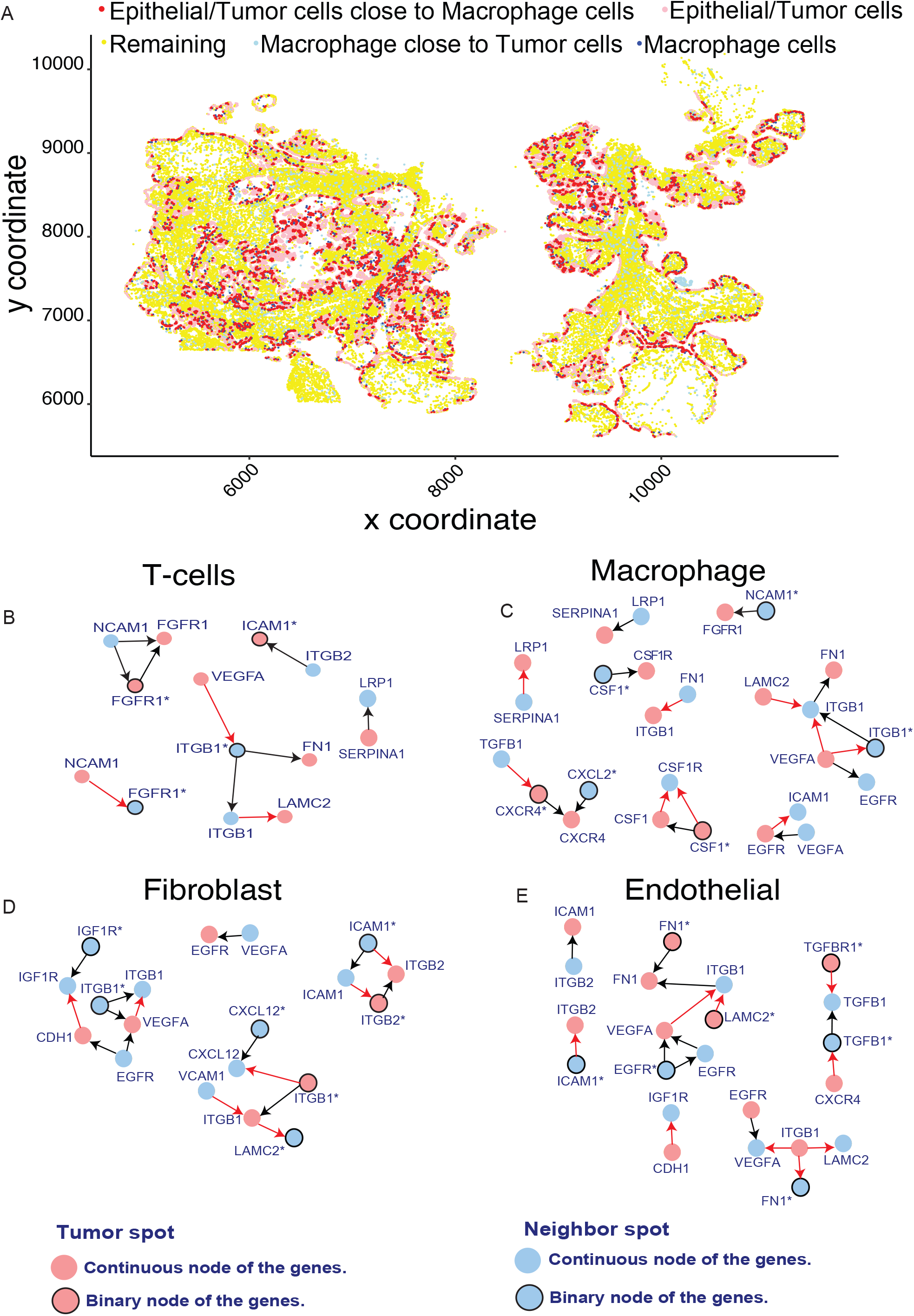

We then assessed the validation of the inferred LR interactions from the 10X data using `LRnetST` in the MERFISH datasets. Specifically, we focused on the 185 reproducible edges, identified in at least two out of the four 10X data sets by `LRnetST`. Among these, only 27 edges had both nodes measured in the MERFISH datasets. Out of these, 19 edges were confirmed in the LR network constructed based on the MERFISH data. The validated edges in `LRnetST`’s result are illustrated in Figure 4A. Specifically, interaction between LRP1 tumor and SERPINA1 stroma was confirmed in multiple MERFISH data sets (Figure 3B-C). Moreover, the bi-directional interaction between ITGB1 tumor — LAMC2 stroma and ITGB1 stroma — VEGFA tumor was also confirmed across multiple tumor slides (Figure 3B-E), further strengthening the confidence of these LR interactions in ovarian tumor tissues.

**Figure 4.**
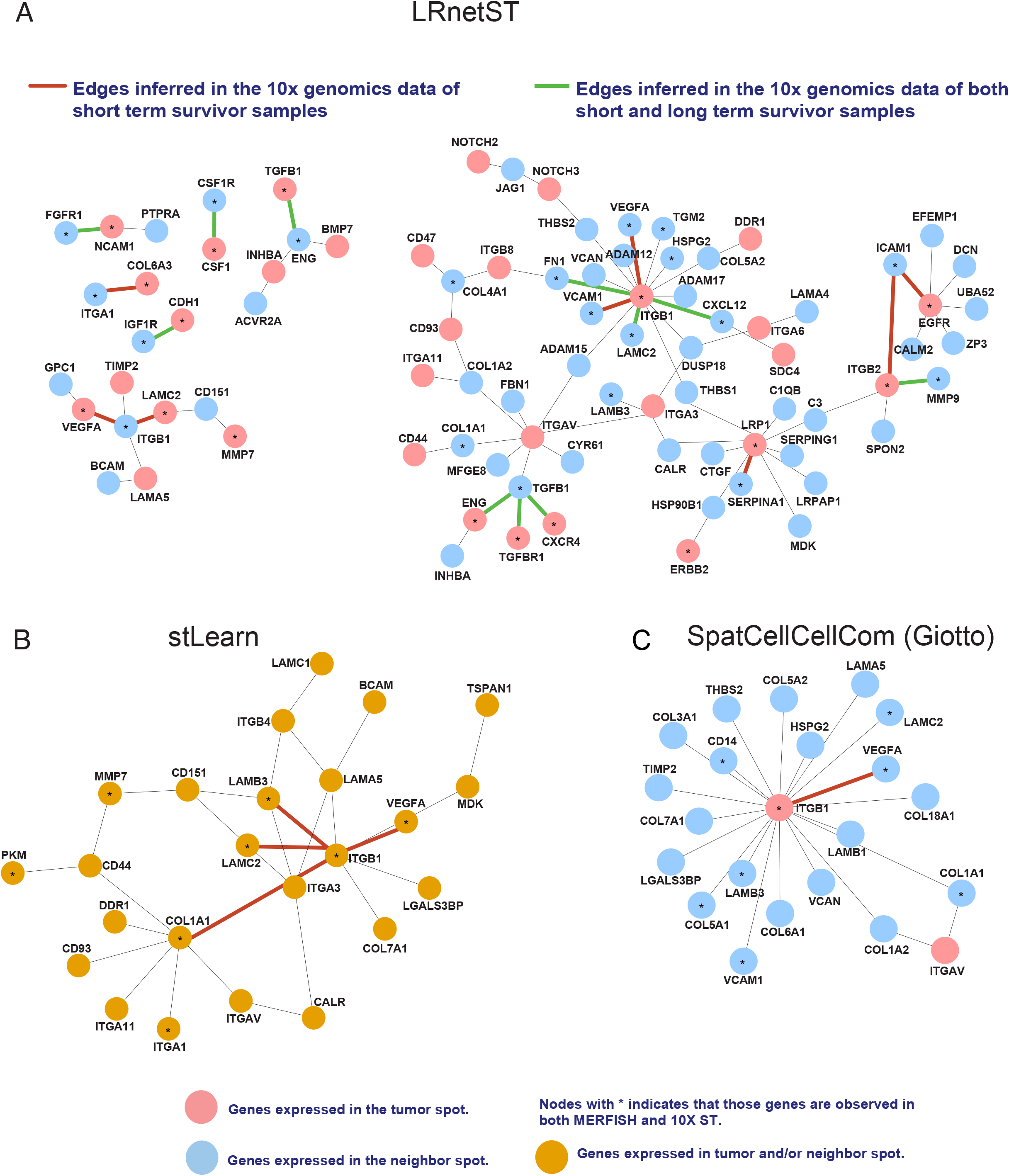

In parallel, we performed the same analysis using `spatCellCellcom` and `stLearn`. As illustrated in Figure 4B-C, only 3 and 1 edges in the results of `stLearn` and `SpatCellCellcom` were validated in the MERFISH data, respectively. Note that the interaction between ITGB1 and VEGFA was inferred by all three methods (Figure 4B-C), suggesting the robust (strong) signal of this LR interaction in ovarian tumor tissues.

## Discussions and conclusions

To gain insights on the molecular basis of tumor micro-environment, in this paper, we introduce a new method — `LRnetST` to infer ligand-receptor interaction networks among adjacent cells using spatial transcriptomic data. `LRnetST` employs novel strategies to address the spatial structure and high dropout rates that are uniquely present in the ST data. We demonstrated the benefit of the strategies of `LRnetST` for modeling the zero-inflated ST data over alternative approaches on simulation examples. We then applied `LRnetST` to 10X ST datasets of four HGSC cancer samples and constructed patient specific ligand-receptor networks. The results reveal both common and distinct ligand-receptor regulations across different patients. A subset of these interactions were validated through a MERFISH data set of independent ovarian tumor samples.

We also compared performance of `LRnetST` and alternative tools on the ovarian ST datasets. The results highlight `LRnetST`’s superior capability in detecting reproducible signals that hold greater biological significance.

In the results of `LRnetST`, LRP1 is identified as a hub node in all four ligand-receptor networks in the 10X data results. LRP1 is reported to make critical contribution to many processes that drive tumorigenesis and tumor progression [46, 47], and has been proposed to be an important diagnostic and prognostic biomarker of epithelial ovarian cancer [42]. Besides the previously reported role of LRP1 in ERK signaling pathway through interaction with MMP9 [42], which was also supported in our results (Figure 2G), we further revealed strong interaction between LRP1 in tumor cells and SERPINA1 (AAT) as well as C1QB in stroma cells (Figure 3B-C, 4A, and S6A, S6C, S6E). The interaction between LRP1 in tumor cells and SERPINA1 in stroma cells was detected in the two short term survivor samples (PFS *<* 6 months) in the 10X data sets, and was consistently confirmed across multiple tumor samples in the MERFISH data (Figure 3B-C, 4A). While the role of LRP1 and SERPINA1 interaction in the context of cancer remains largely unexplored, existing research in cardiovascular studies has highlighted its significance in an anti-inflammatory mechanism [48]. By analogy, one may hypothesize that the interaction between LRP1 in tumor cells and SERPINA1 in stroma cells could contribute to an immunosuppressive effect within the tumor microenvironment.

Interaction between LRP1 and C1QB, on the other hand, was only detected in the two long term survivor samples in 10X data sets. C1QB is a pattern recognition molecule of innate immunity and can be locally synthesized by macrophages and dendritic cells [34]. The connection between LRP1 and C1QB detected in the two chemo-sensitive patients suggests a potential role of LRP1-C1QB crosstalk network in modulating immune response which may then contribute to the chemo-treatment effectiveness in HGSC patients. Unfortunately, C1QB was not detected in the MERFISH data sets, so validation of this interaction using additional resources is warranted as future studies.

Another hub node detected in all four tumors based on 10X data sets was ITGB1, which has been inferred to interact with VEGFA and VCAM1 in the two chemo-refractory patients. These interactions were also validated on the MERFISH data. Intercellular communication mediated via interaction between VEGFA and ITGB1 has been reported to contribute to the precise coordination of multiple biological processes, including development, differentiation, and inflammation [44]. Recently, interactions through VEGFA and ITGB1 was reported to underlie the crosstalk between epithelial cells and fibroblasts relating to tumor inflammation in Pancreatic ductal cancer [49]. ITGB1 and VCAM1 are both cell-adhesion molecules that influence immune response and anti-immunity. VCAM1 mediates intercellular adhesion by specific binding to the integrin ITGB1 on leukocytes, and thus this interaction plays a pathophysiologic role in leukocyte emigration to sites of inflammation [45]. The interaction between ITGB1 and VCAM1 has also been reported in the context of cell migration between brain endothelial cells (BECs) and central memory T cells (Tcm) [50]. However, our analysis, for the first time, suggests the potential roles of the cross-talk between ITGB1, VEGFA and/or VCAM1 among ovarian cancer patients.

In conclusion, spatial transcriptomic data offers an unprecedented opportunity to unravel the molecular mechanisms underlying cell-cell interactions within the tumor micro - environment. The `LRnetST` method, proposed in this study, proves to be a valuable tool for constructing ligand-receptor interaction networks based on spatial transcriptomic (ST) data. Similar to other high-dimensional network construction methods, the performance of `LRnetST` may be impacted by the dimension of the gene space, as demonstrated in our simulation study. This underscores the significance of leveraging prior domain knowledge to constrain the Directed Acyclic Graph (DAG) search space when applying `LRnetST`, as exemplified in our ovarian cancer ST data analysis through the use of existing ligand-receptor networks. The outcomes of these enhanced analyses promise to unveil crucial contributors driving cell-cell interactions, potentially leading to the identification of new predictive biomarkers or therapeutic targets. An R package of `LRnetST` is made available as a github repository https://github.com/jie108/LRnetST.

## Ethics approval and consent to participate

Not applicable.

## Author contributions statement

Writing: SC, PY, WW, JP and PW. Data analysis and Visualization: SC, JP, PW, WW, SFB, PY, APC and SCM. Result interpretation: SC, JP, PW, WW, SFB, PY, APC and SCM. Supervision: WW, SCM, JP and PW. Funding Acquisition: WW and PW.

## Acknowledgments

This work is supported by grants (U24CA271114, U01CA271407) and a grant (R01CA268380) from the National Cancer Institute. We also thank Dr. Guocheng Yuan and his team members for their instruction on the use of Giotto.

## Supplementary Information

**Additional file 1** contains supplementary methods description and tables.

## Availability of data and materials

An R package of `LRnetST` is made available as a github repository https://github.com/jie108/LRnetST. All the codes along with the data will be made available through this github repository.

---

**Figure S1.**
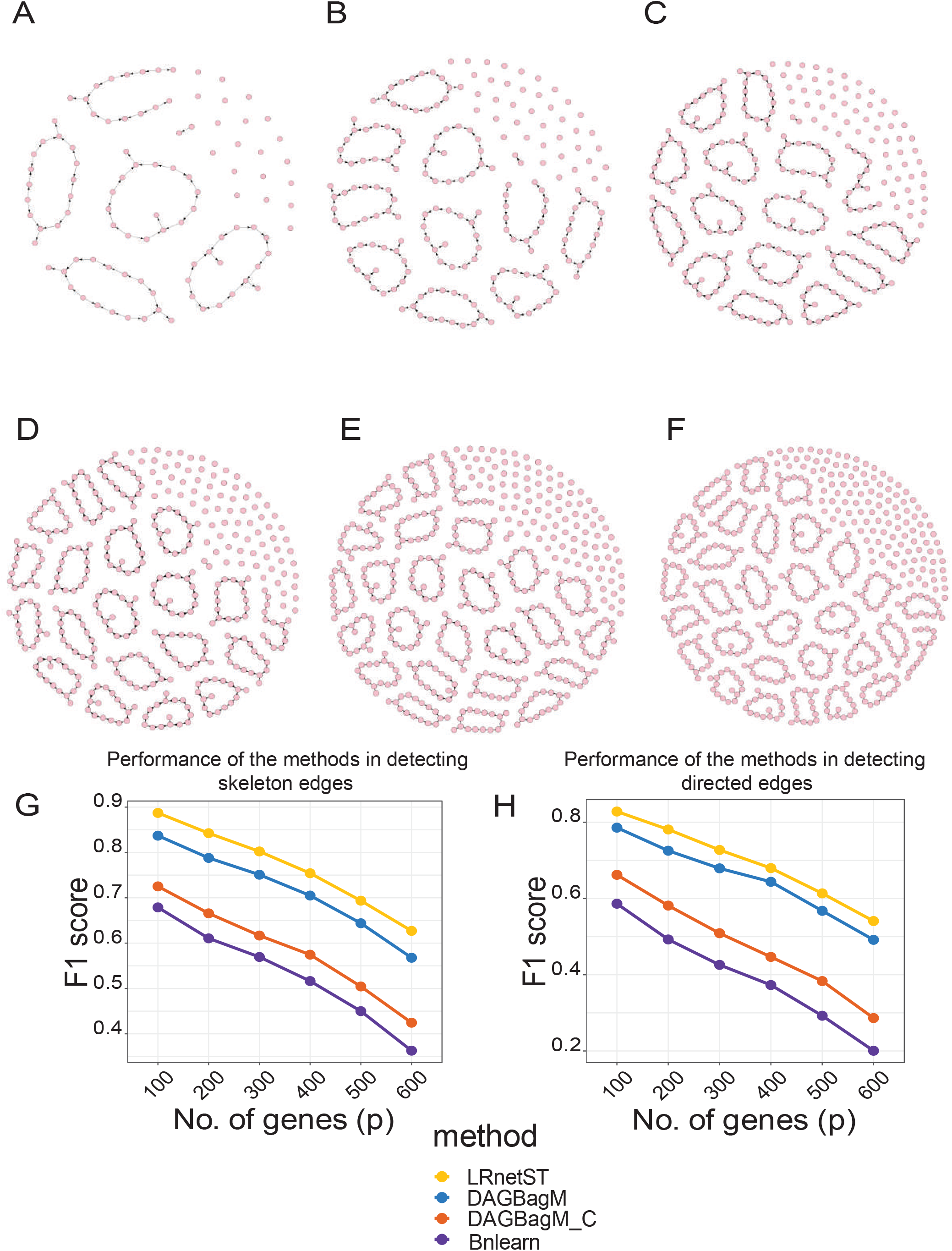

**Figure S2.**
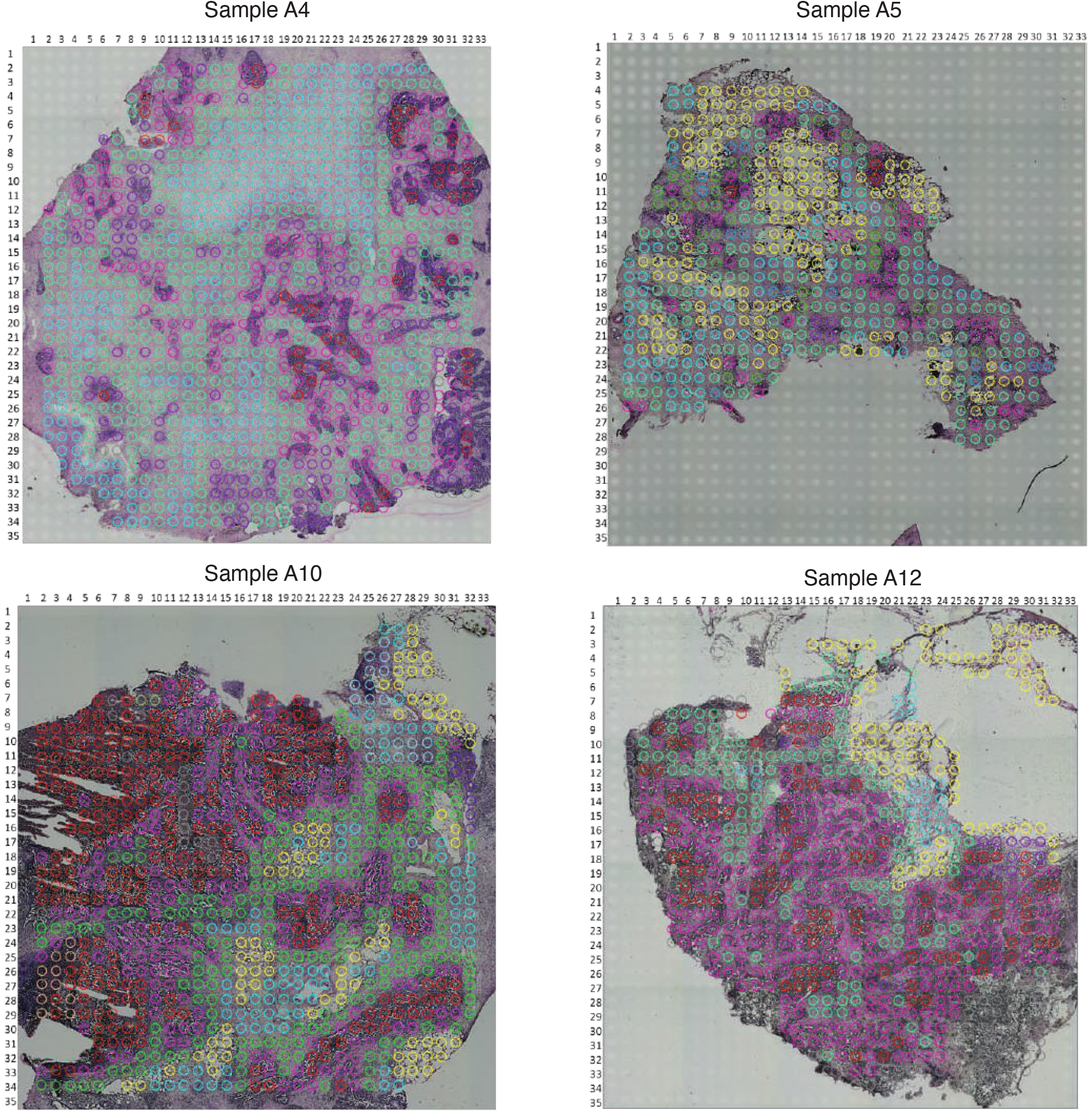

**Figure S3.**
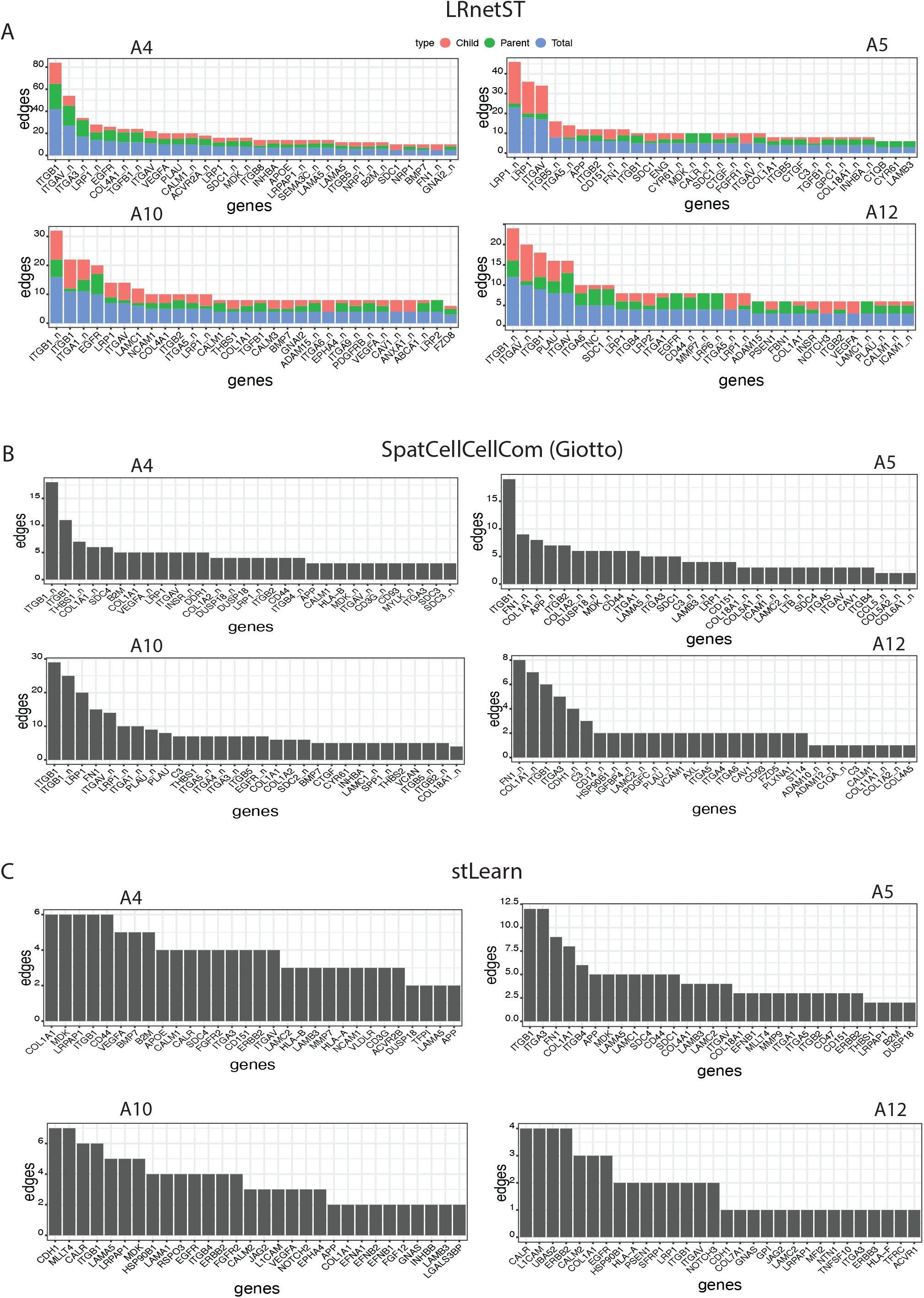

**Figure S4.**
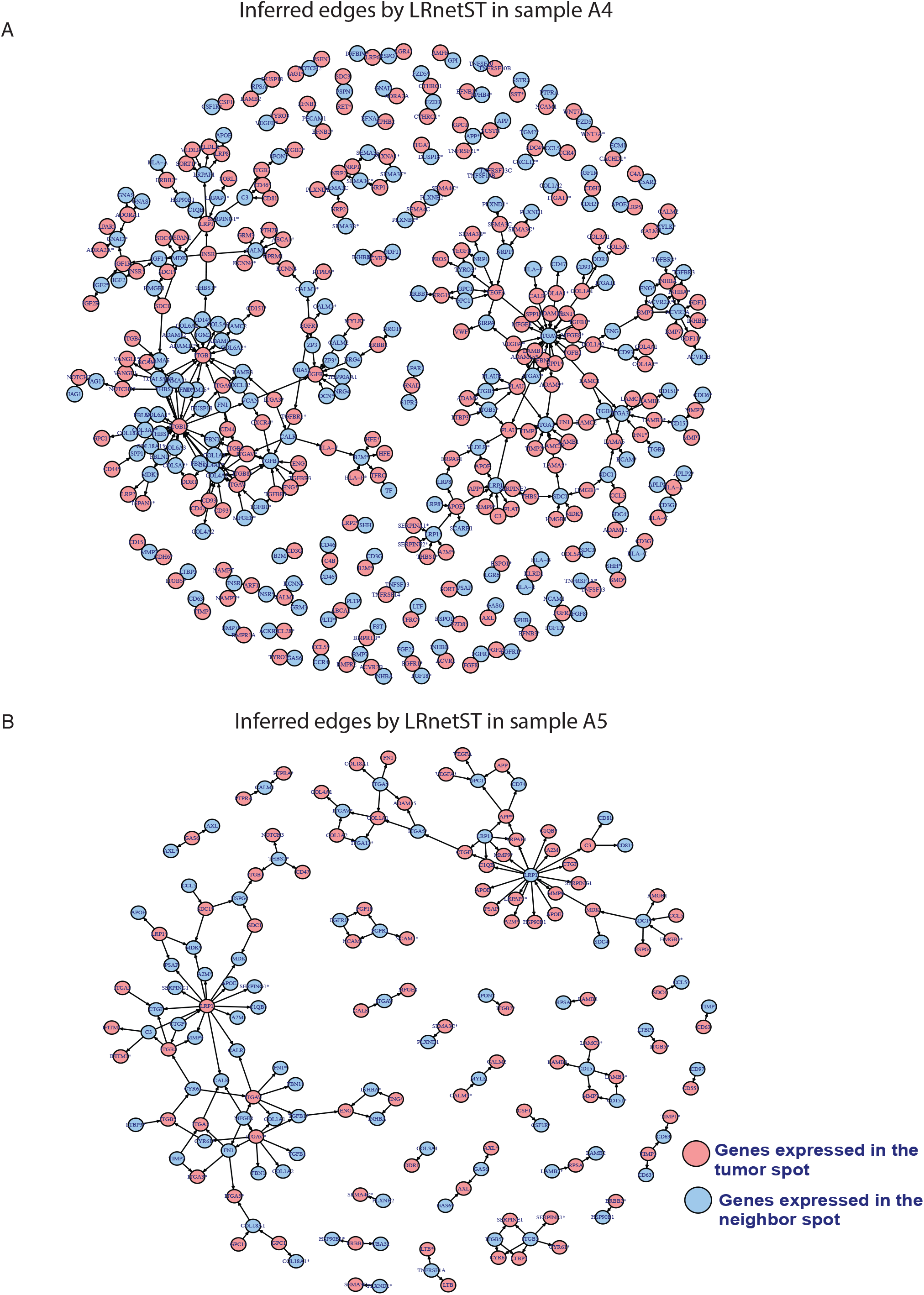

**Figure S5.**
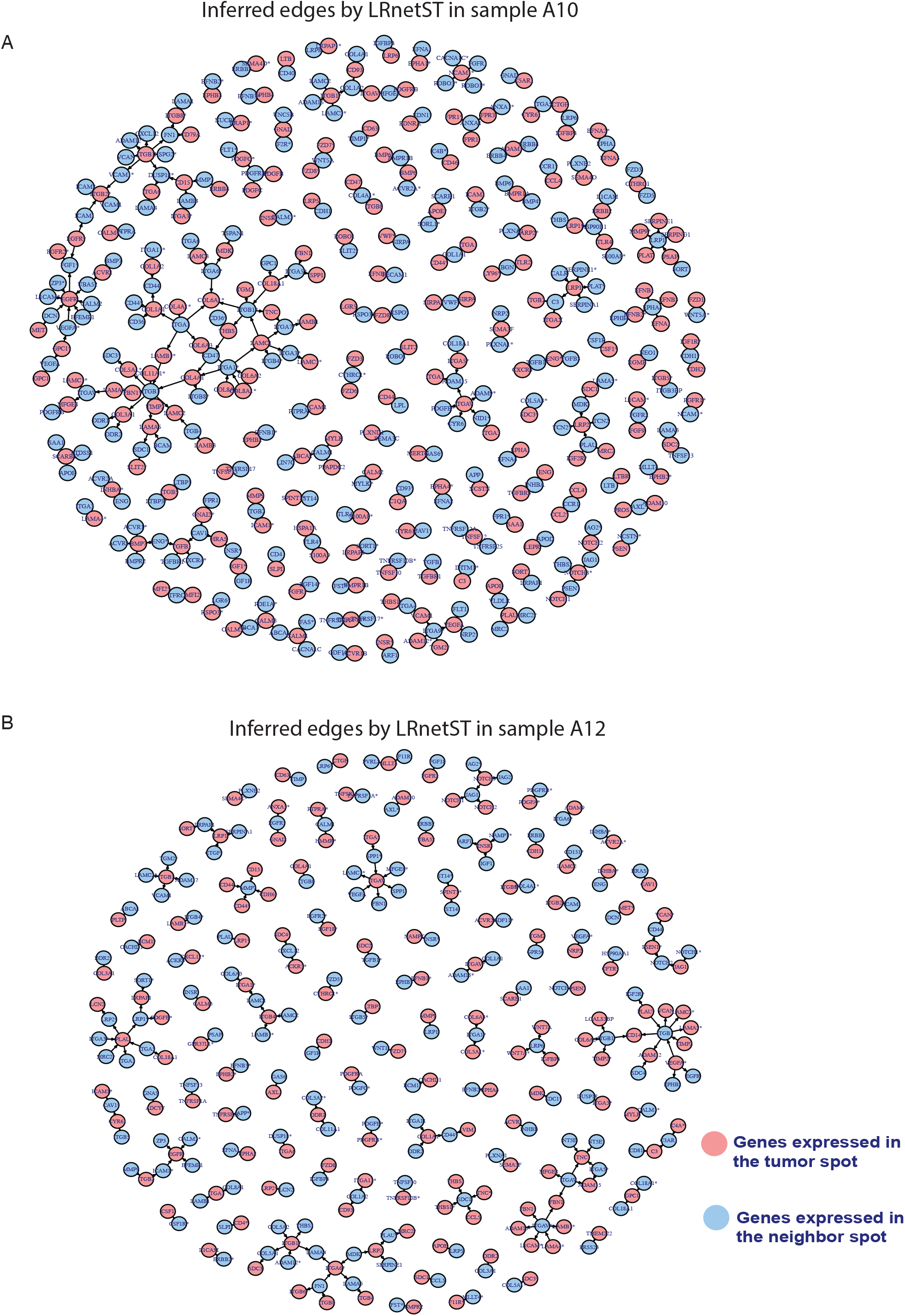

**Figure S6.**
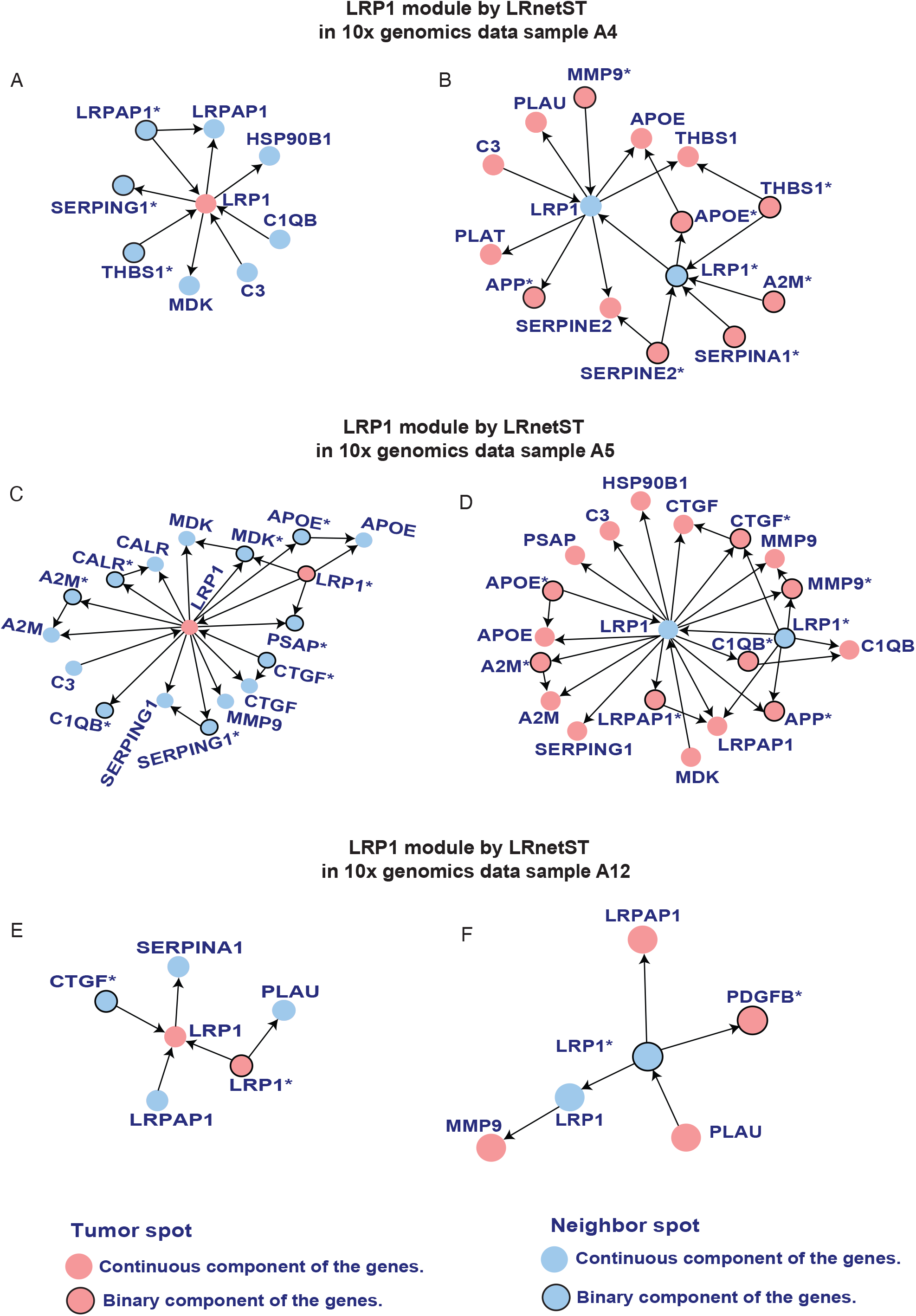

**Figure S7.**
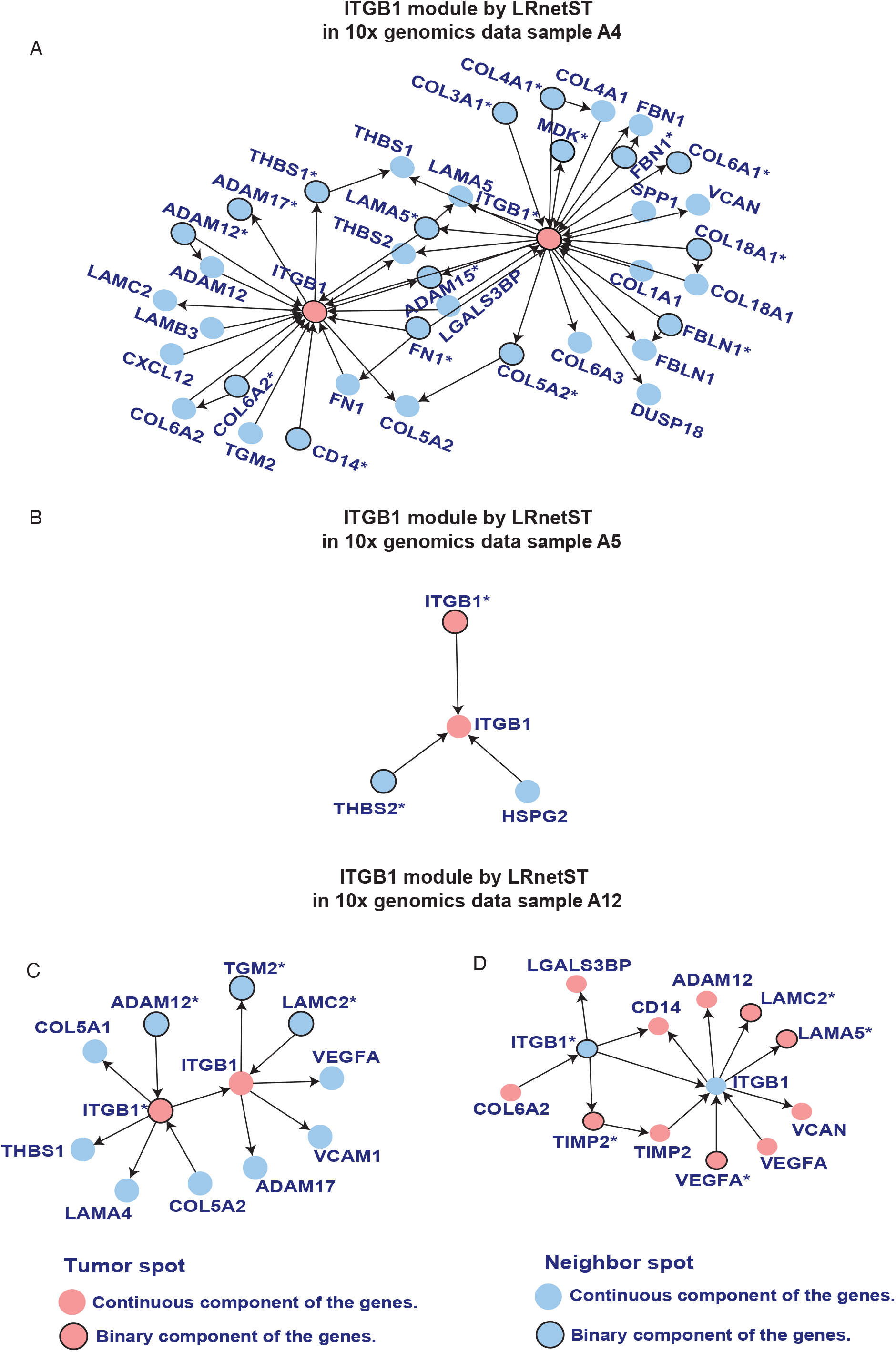

**Figure S8.**
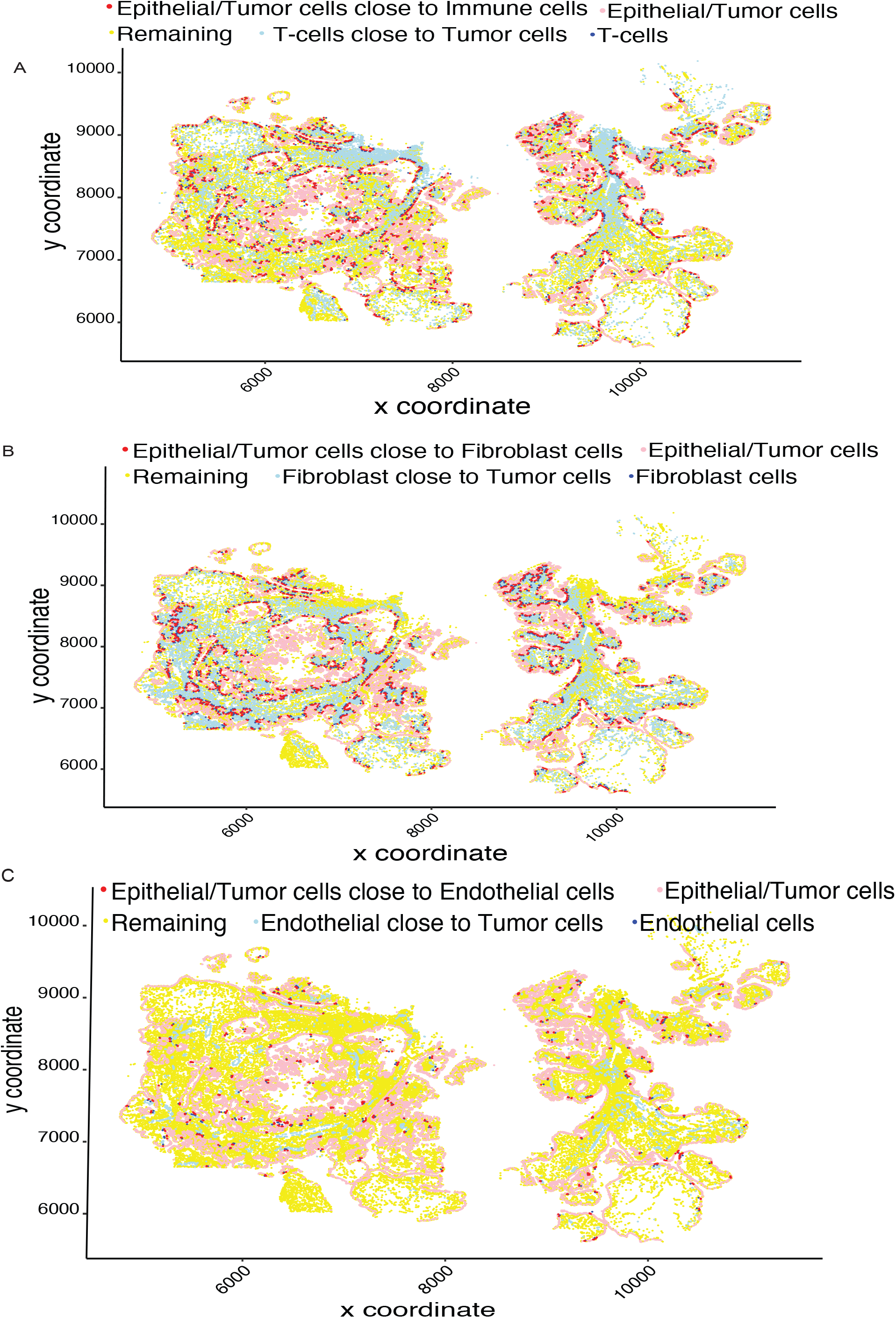

**Figure S9.**
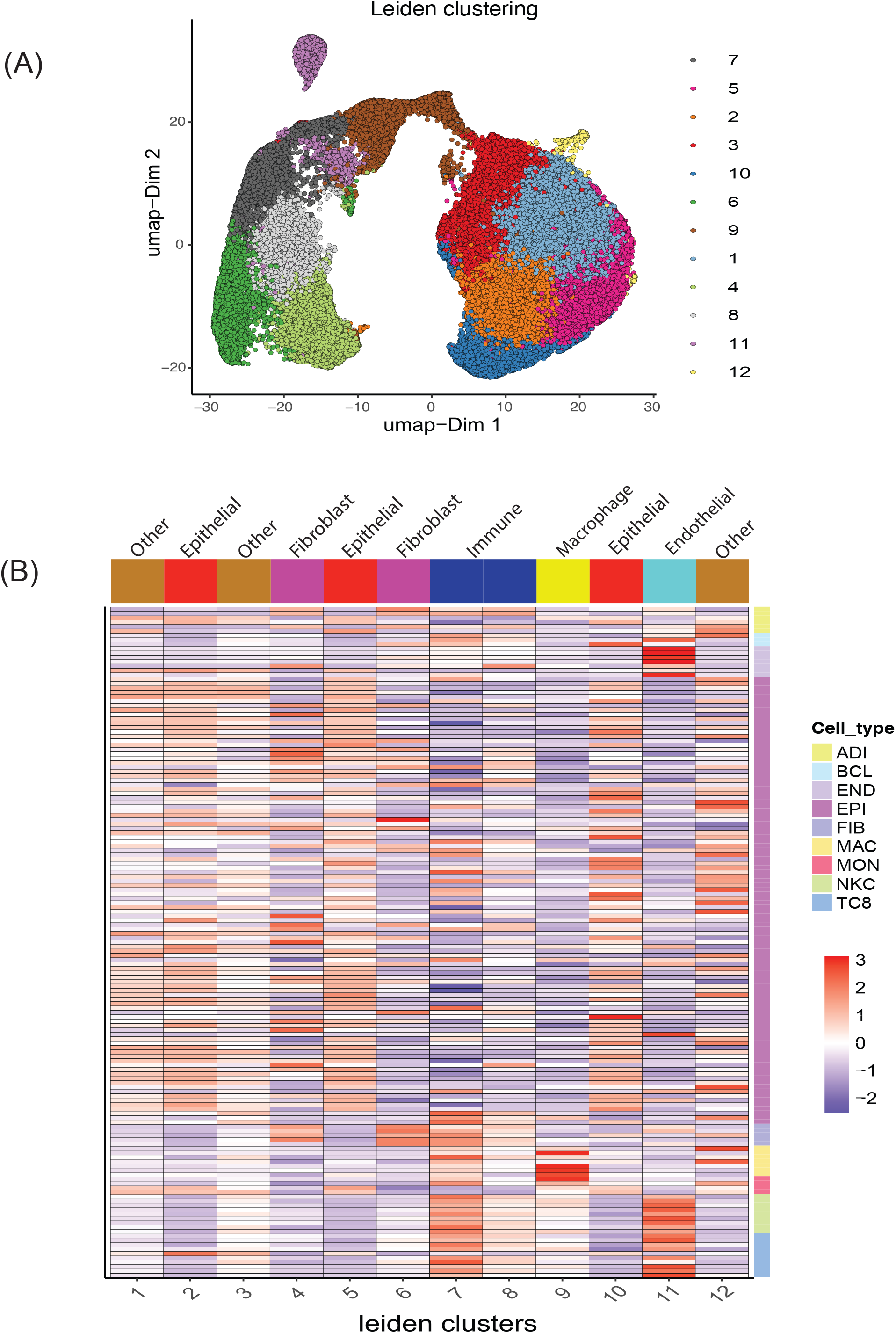

## References

1. Cancer Genome Atlas Research Network et al. Integrated genomic analyses of ovarian carcinoma. Nature, 474:609– 615, 2011.

2. A. M. Patch et al. Whole-genome characterization of chemo-resistant ovarian cancer. Nature, 521(7553):489–494, 2015.

3. R. W. Tothill et al. Novel molecular subtypes of serous and endometrioid ovarian cancer linked to clinical outcome. Clin. Cancer Res., 14:5198–5208, 2008.

4. H. Zhang et al. Integrated proteogenomic characterization of human high-grade serous ovarian cancer. Cell, 166(3):755–765, 2016.

5. S. Chowdhury, J.J. Kennedy, I.G. Ivey, O.D. Murillo, N. Hosseini, et al. Proteogenomic analysis of chemorefractory high-grade serous ovarian cancer. Cell, 186(16):3476–3498, 2023.

6. B. Zhang et al. Revisiting ovarian cancer microenvironment: a friend or a foe? Protein Cell, 9(8):674–692, 2018.

7. T. Tsujikawa, J. Mitsuda, H. Ogi, A. Miyagawa-Hayashino, E. Konishi, K. Itoh, and S. Hirano. Prognostic significance of spatial immune profiles in human solid cancers. Cancer Sci, 111(10):3426–3434, 2020.

8. M. Efremova, M. Vento-Tormo, S. A. Teichmann, and R. Vento-Tormo. Cellphonedb: inferring cell–cell communication from combined expression of multi-subunit ligand–receptor complexes. Nature protocols, 15(4):1484–1506, 2020.

9. S. Cabello-Aguilar et al. Singlecellsignalr: inference of intercellular networks from single-cell transcriptomics. Nucleic Acids Research, 48(10), 2020.

10. S. Jin et al. Inference and analysis of cell-cell communication using cellchat. Nature communications, 12(1):1–20, 2021.

11. F. Noel et al. Dissection of intercellular communication using the transcriptome-based framework icellnet. Nature communications, 12(1089), 2021.

12. J. S. Nagai et al. Crosstalker: analysis and visualization of ligand–receptor networks. Bioinformatics, 37(22):4263–4265, 2021.

13. R. Browaeys, W. Saelens, and W. Saeys. Nichenet: modeling intercellular communication by linking ligands to target genes. Nature methods, 17(2):159–162, 2020.

14. J. Cheng, J. Zhang, Z. Wu, and X. Sun. Inferring microenvironmental regulation of gene expression from single-cell rna sequencing data using scmlnet with an application to covid-19. Briefings in bioinformatics, 22(2):988–1005, 2021.

15. J. Cheng, J. Zhang, Z. Wu, and X. Sun. Cytotalk: De novo construction of signal transduction networks using single-cell transcriptomic data. Science Advances, 7(16):eabf1356, 2021.

16. M. Asp, J. Bergenstråhle, and J. Lundeberg. Spatially resolved transcriptomes—next generation tools for tissue exploration. BioEssays, 42(10), 2020.

17. V. Marx. Method of the year: spatially resolved transcriptomics. Nature Methods, 18(1):9–14, 2021.

18. P.L. Ståhl et al. Visualization and analysis of gene expression in tissue sections by spatial transcriptomics. Science, 353(6294):78–82, 2016.

19. R. Dries, Q. Zhu, R. Dong, C. H. L. Eng, H. Li, K. Liu, et al. Giotto: a toolbox for integrative analysis and visualization of spatial expression data. Genome biology, 22:1–31, 2021.

20. D. Pham, X. Tan, J. Xu, L. F. Grice, P. Y. Lam, et al. stlearn: integrating spatial location, tissue morphology and gene expression to find cell types, cell-cell interactions and spatial trajectories within undissociated tissues. BioRxiv, 22:2020–2025, 2020.

21. T. Verma and J. Pearl. Equivalence and synthesis of causal models. In Henrion, M., Shachter, R. Kanal, L., and Lemmer, J., editors, Proceeding of the Sixth Conference on Uncertainty in Artificial Intelligence, pages 220–227, 1991.

22. D. Geiger and D. Heckerman. Learning gaussian networks. In Proceedings of the Tenth international conference on Uncertainty in artificial intelligence, pages 235–243. Morgan Kaufmann Publishers Inc., 1994.

23. P. Spirtes, C. Glymour, and R. Scheines. Causation, prediction, and search, volume 81. MIT press, 2001.

24. M. Kalisch and P. Bühlmann. Estimating high-dimensional directed acyclic graphs with the pc-algorithm. The Journal of Machine Learning Research, 8:613–636, 2007.

25. I. Tsamardinos, L.E. Brown, and C.F. Aliferis. The max-min hill-climbing bayesian network structure learning algorithm. Machine learning, 65(1):31–78, 2006.

26. S. Chowdhury, R. Wang, Q. Yu, C. J. Huntoon, L. M. Karnitz, S. H. Kaufmann, and et al. Dagbagm: learning directed acyclic graphs of mixed variables with an application to identify protein biomarkers for treatment response in ovarian cancer. BMC Bioinformatics, 23, 2022.

27. M. Scutari. Learning bayesian networks with the bnlearn r package. Journal of Statistical Software, 35(3), 2010.

28. S. Ferri-Borgogno, Y. Zhu, J. Sheng, J. K. Burks, J. A. Gomez, K. K. Wong, et al. Spatial transcriptomics depict ligand–receptor cross-talk heterogeneity at the tumor-stroma interface in long-term ovarian cancer survivors. Cancer Research, 83(9):1503–1516, 2023.

29. J. F. Navarro et al. St pipeline: an automated pipeline for spatial mapping of unique transcripts. Bioinformatics, 33(16):2591–2593, 2017.

30. A. Dobin et al. Star: ultrafast universal rna-seq aligner. Bioinformatics, 29(1):15–21, 2013.

31. L. Geistlinger, S. Oh, M. Ramos, L. Schiffer, R. S. LaRue, C. M. Henzler, and et al. Multiomic analysis of subtype evolution and heterogeneity in high-grade serous ovarian carcinoma. Cancer research, 80(20):4335–4345, 2020.

32. Vizgen merfish ffpe human immuno-oncology data set. Vizgen, 2022.

33. J. Pearl. Causality: models, reasoning and inference, volume 29. Cambridge Univ Press, 2000.

34. Jun Zhu, Bin Zhang, Erin N Smith, Becky Drees, Rachel B Brem, Leonid Kruglyak, Roger E Bumgarner, and Eric E Schadt. Integrating large-scale functional genomic data to dissect the complexity of yeast regulatory networks. Nature genetics, 40(7):854–861, 2008.

35. J. Zhu, P. Sova, Q. Xu, K. M. Dombek, E. Y. Xu, and H. et al Vu. Stitching together multiple data dimensions reveals interacting metabolomic and transcriptomic networks that modulate cell regulation. PLoS Biology, 10(4), 2012.

36. W. H. Sung, C. Gong, C. Myun-Seok, and Z. Hua. Estimation of directed acyclic graphs through two-stage adaptive lasso for gene network inference. Journal of the American Statistical Association, 111(515):1004–1019, 2016.

37. N. Friedman, M. Linial, I. Nachman, and D. Pe’er. Using bayesian networks to analyze expression data. Journal of computational biology, 7(3-4):601–620, 2000.

38. D. PeÕer, A. Regev, G. Elidan, and N. Friedman. Inferring subnetworks from perturbed expression profiles. Bioinformatics, 17(suppl 1):S215–S224, 2001.

39. K. Sachs, O. Perez, D. Pe’er, D.A. Lauffenburger, and G.P. Nolan. Causal protein-signaling networks derived from multiparameter single-cell data. Science Signalling, 308(5721):523, 2005.

40. J. Ramilowski, T. Goldberg, J. Harshbarger, et al. A draft network of ligand–receptor-mediated multicellular signalling in human. Nat Commun, 6(7866):755–765, 2015.

41. X. He and J. Zhang. Why do hubs tend to be essential in protein networks? PLoS genetics, 2(6), 2006.

42. W. Zhou, J. Ma, H. Zhao, Q. Wang, X. Guo, L. Chen, and et al. Serum exosomes from epithelial ovarian cancer patients contain lrp1, which promotes the migration of epithelial ovarian cancer cell. Molecular Cellular Proteomics, 22(4), 2023.

43. D. K. Strickland, S. C. Muratoglu, and T. M. Antalis. Serpin–enzyme receptors: Ldl receptor-related protein 1. Methods in enzymology, 499:17–31, 2011.

44. R. Miao, X. Dong, J. Gong, Y. Li, X. Guo, and J. et al. Wang. Examining the development of chronic thromboembolic pulmonary hypertension at the single-cell level. Hypertension, 79(3):562–574, 2022.

45. J. Wu, Z. P. Chen, A. Q. Shang, W. W. Wang, Z. N. Chen, Y. J. Tao, and et al. Systemic bioinformatics analysis of recurrent aphthous stomatitis gene expression profiles. Oncotarget, 8(67), 2017.

46. A. P. Lillis, L. B. Van Duyn, J. E. Murphy-Ullrich, and D. K. Strickland. Ldl receptor-related protein 1: unique tissue-specific functions revealed by selective gene knockout studies. Physiological reviews, 88(3):887–918, 2008.

47. P. Xing, Z. Liao, Z. Ren, J. Zhao, F. Song, G. Wang, and et al. Roles of low-density lipoprotein receptor-related protein 1 in tumors. Chinese journal of cancer, 35(1):1–8, 2016.

48. N. Potere, M. G. Del Buono, A. G. Mauro, A. Abbate, and S. Toldo. Low density lipoprotein receptor-related protein-1 in cardiac inflammation and infarct healing. Frontiers in cardiovascular medicine, 6(51), 2019.

49. B. Kinny-Köster, S. Guinn, J. A. Tandurella, J. T. Mitchell, D. N. Sidiropoulos, M. Loth, and et al. Inflammatory signaling in pancreatic cancer transfers between a single-cell rna sequencing atlas and co-culture. bioRxiv, 2022.

50. X. Zhang, R. Wang, H. Chen, C. Jin, Z. Jin, J. Lu, and et al. Aged microglia promote peripheral t cell infiltration by reprogramming the microenvironment of neurogenic niches. Immunity Ageing, 19(1), 2022.

