## Additional file for "Learning directed acyclic graphs for ligands and receptors based on spatially resolved transcriptomic analysis of ovarian cancer"

### ADDITIONAL FILE: SUPPLEMENTARY MATERIAL

#### A.1 `LRnetST` pipeline

Consider the ST data set of one biology sample (one ST slice). Refer to the ST spots(grids) enriched of tumor cells as *Index* Spots, while their closest spots that are enriched of non-tumor cells as *Neighbor* Spots. Below we outline the details of the `LRnetST` algorithm:

`LRnetST`: Directed acyclic graph learning through bootstrap aggregation for ST data.

1. Define Index-Neighbor-Spot pairs: For each *Index* Spot, identify its closest *Neighbor* Spot according to the Euclidean distances between spot locations on the ST slice. Filter away the Index-Neighbor-Spot pairs if the distances are more than a certain cutoff (e.g. 2×the grid unit on the ST slice). Denote the number of the remaining Index-Neighbor-Spot pairs as $n$.
2. Generate down-sampled data sets $D_b$ for $b = 1, \cdots, B$: Denote $N$ as the median of the total UMI counts of samples (ST spots) in the original gene expression data. Use $N$ as the maximum total UMI count value to perform the downsampling normalization and generate $B$ downsampled data $\{D_b\}_{b=1}^B$.
3. For each downsampled data $D_b$ ($b = 1, \cdots, B$):

   Gene filter: Filter genes with more than 50 % missing across all the samples (spots). Further take the overlap of the resulting set and the ligand-receptor database. Denote the size of the final gene set as $p_b$. For simplification of the notation, we just use $p$ in below.

   Generate the $b^{th}$ Neighbor Integrated Matrix $NIM_b$ and its bootstrap re-sample $M_b$:

   a. Stack the gene expressions of the $p$ genes in the $n$ Index Spots (a sub-matrix of $D_b$) with that of their matching Neighbor Spots (another sub-matrix of $D_b$). The resulting matrix, $Initial - NIM_b$, has a dimension of $2p \times n$.

   b. Derive $NIM_b$ by adding $2p$ binary variables to $Initial - NIM_b$, one for each of the $2p$ gene expression features in $Initial - NIM_b$. The binary variables take 1 or 0 in a sample depending on whether the corresponding gene expressions are non-zero or zero in that sample. Moreover, convert the expression features according to the binary node: if the binary node takes value 1, then keep the original value; otherwise, replace the original value with $NA$.

   c. A bootstrap sample (with the same number of columns) of $NIM_b$ is generated by sampling the columns with replacement and is denoted as $M_b$, which has a dimension of $4p \times n$.

   Ensemble learning: For each $M_b$, learn a DAG $\mathcal{G}_b$ as follows ($b = 1, \cdots, B$): .

   Initial step: Start with an empty graph $\mathcal{G}^0$, and calculate the initial score $S(\mathcal{G}^0, M_b)$. The set of possible operations ($O^0$) is set to include additions of all possible edges.

   Updating step: In the $t^{th}$ step ($t \geq 1$), denote the current graph by $\mathcal{G}^t$, the current score by $S(\mathcal{G}^t, M_b)$ and the current set of eligible operations by $O^t$ (which includes any addition, deletion or reversal of an edge without violating acyclicity or whitelist).

   a. For every eligible operation $O \in O^t$, calculate the score change: $\epsilon^t = S(O(\mathcal{G}^t), M_b) - S(\mathcal{G}^t, M_b)$, where $O(\mathcal{G}^t)$ denotes the graph after applying the operation $O$ on $\mathcal{G}^t$.

   b. Calculate the minimum score change $\epsilon^t_{min} = min_{O \in O^t} \epsilon^t$ and denote the corresponding operation as $\tilde{O}$.

   c. Updating/Stopping rule: If $\epsilon^t_{min} \geq 0$, stop and take $\mathcal{G}_b = \mathcal{G}^t$ as the estimated graph on $b^{th}$ downsampled data; else set $\mathcal{G}^{t+1} = O(\mathcal{G}^t)$ and $t \leftarrow t + 1$ and repeat the updating step.

   Aggregation: $\mathcal{G}^* = \text{argmin}_{\mathcal{G} \in \mathbb{G}(\mathbb{N})} score_d(\mathcal{G} : \mathbb{G}^e)$, where $\mathbb{G}(\mathbb{N})$ is the DAG space with the node set $\mathbb{N}$, and $\mathbb{G}^e = \{\mathcal{G}_b,\ b = 1, \cdots, B\}$, $score_d(\mathcal{G} : \mathbb{G}^e) = \frac{1}{B} \sum_{b=1}^B d(\mathcal{G}, \mathcal{G}_b)$, and $d(\cdot, \cdot)$ is the structural Hamming distance on $\mathbb{G}(\mathbb{N})$.

#### A.2 Simulation experiments: additional details

Table A.1: Simulation results for different combinations of the number of genes ($p$) and the number of edges ($||E||$). The range of signal-to-noise-ratio (SNR) is set to be $[0.5, 1.5]$ and the sample sizes are $n = 200, 400$, respectively. The reported numbers are averaged over 100 independent replicates.

| Method | No. of genes (p) | Sample size (n) | Detecting skeleton edges | | Detecting directed edges | |
|---|---|---|---|---|---|---|
| | | | Power (TPR) | FDR | Power (TPR) | FDR |
| LRnetST | 100 | 400 | 0.84 | 0.06 | 0.76 | 0.09 |
| DAGBagM | | | 0.75 | 0.13 | 0.68 | 0.15 |
| DAGBagMC | | | 0.65 | 0.18 | 0.56 | 0.19 |
| bnlearn | | | 0.62 | 0.25 | 0.49 | 0.27 |
| LRnetST | 200 | 400 | 0.77 | 0.07 | 0.69 | 0.1 |
| DAGBagM | | | 0.68 | 0.15 | 0.60 | 0.175 |
| DAGBagMC | | | 0.57 | 0.20 | 0.46 | 0.21 |
| bnlearn | | | 0.53 | 0.28 | 0.38 | 0.30 |
| LRnetST | 300 | 400 | 0.71 | 0.078 | 0.62 | 0.12 |
| DAGBagM | | | 0.63 | 0.166 | 0.54 | 0.19 |
| DAGBagMC | | | 0.51 | 0.22 | 0.38 | 0.23 |
| bnlearn | | | 0.48 | 0.30 | 0.31 | 0.32 |
| LRnetST | 400 | 400 | 0.64 | 0.082 | 0.56 | 0.135 |
| DAGBagM | | | 0.57 | 0.18 | 0.50 | 0.2 |
| DAGBagMC | | | 0.46 | 0.235 | 0.32 | 0.26 |
| bnlearn | | | 0.43 | 0.33 | 0.26 | 0.34 |
| LRnetST | 500 | 400 | 0.56 | 0.089 | 0.48 | 0.15 |
| DAGBagM | | | 0.49 | 0.2 | 0.41 | 0.23 |
| DAGBagMC | | | 0.38 | 0.25 | 0.27 | 0.27 |
| bnlearn | | | 0.35 | 0.37 | 0.19 | 0.365 |
| LRnetST | 600 | 400 | 0.48 | 0.096 | 0.4 | 0.165 |
| DAGBagM | | | 0.40 | 0.23 | 0.32 | 0.26 |
| DAGBagMC | | | 0.30 | 0.275 | 0.18 | 0.30 |
| bnlearn | | | 0.27 | 0.40 | 0.12 | 0.39 |
| LRnetST | 100 | 200 | 0.72 | 0.07 | 0.67 | 0.1 |
| DAGBagM | | | 0.65 | 0.14 | 0.60 | 0.166 |
| DAGBagMC | | | 0.56 | 0.20 | 0.50 | 0.22 |
| bnlearn | | | 0.53 | 0.28 | 0.45 | 0.30 |
| LRnetST | 200 | 200 | 0.69 | 0.079 | 0.61 | 0.11 |
| DAGBagM | | | 0.60 | 0.16 | 0.54 | 0.19 |
| DAGBagMC | | | 0.48 | 0.24 | 0.43 | 0.25 |
| bnlearn | | | 0.45 | 0.34 | 0.39 | 0.34 |
| LRnetST | 300 | 200 | 0.62 | 0.085 | 0.56 | 0.135 |
| DAGBagM | | | 0.56 | 0.185 | 0.49 | 0.21 |
| DAGBagMC | | | 0.39 | 0.26 | 0.36 | 0.27 |
| bnlearn | | | 0.37 | 0.37 | 0.32 | 0.37 |
| LRnetST | 400 | 200 | 0.57 | 0.091 | 0.48 | 0.15 |
| DAGBagM | | | 0.50 | 0.21 | 0.42 | 0.22 |
| DAGBagMC | | | 0.31 | 0.29 | 0.28 | 0.30 |
| bnlearn | | | 0.28 | 0.40 | 0.23 | 0.41 |
| LRnetST | 500 | 200 | 0.48 | 0.097 | 0.40 | 0.16 |
| DAGBagM | | | 0.40 | 0.24 | 0.33 | 0.25 |
| DAGBagMC | | | 0.23 | 0.32 | 0.21 | 0.33 |
| bnlearn | | | 0.21 | 0.42 | 0.16 | 0.43 |
| LRnetST | 600 | 200 | 0.41 | 0.1 | 0.32 | 0.18 |
| DAGBagM | | | 0.33 | 0.27 | 0.24 | 0.30 |
| DAGBagMC | | | 0.17 | 0.35 | 0.13 | 0.37 |
| bnlearn | | | 0.15 | 0.46 | 0.08 | 0.48 |

#### A.3 Application to ovarian cancer ST data

Table A.2: Numbers of spots enriched of tumor and/or stromal cells, and number ofligand/receptor genes in 10X ST data sets offour tumors.

| Tumor sample ID | PFS | # Spots (Tumor and Tumor/Stromal) | # Spots labelled as Stromal | # Adjacent tumor-stromal spot pairs (sample size) | # Ligand and receptor genes |
|---|---|---|---|---|---|
| A4 | $>$ 6 months | 240 | 350 | 202 | 184 |
| A5 | $>$ 6 months | 51 | 147 | 51 | 111 |
| A10 | $<$ 6 months | 424 | 183 | 238 | 230 |
| A12 | $<$ 6 months | 359 | 118 | 236 | 196 |

Table A.3: Numbers of tumor and immune/stromal cells and number ofligand/receptor genes in MERFISH data sets offour tumor slices

| MERFISH sample | Stromal cell type | # Tumor cells | # Adjacent tumor-stromal cells | # Ligand and receptor genes |
|---|---|---|---|---|
| Patient 1 | Macrophage | 46377 | 18470 | 384 |
| | T-cells | 7045 | 3071 | 222 |
| | Endothelial | 8983 | 3879 | 448 |
| Patient 2 slice 1 | Macrophage | 21249 | 7817 | 350 |
| | Fibroblast | 6806 | 3496 | 366 |
| | Endothelial | 3427 | 1324 | 444 |
| Patient 2 slice 2 | Macrophage | 5300 | 2310 | 406 |
| | T-cells | 2308 | 1229 | 260 |
| | Endothelial | 446 | 221 | 448 |
| | Fibroblast | 2313 | 1403 | 360 |
| Patient 2 slice 3 | Fibroblast | 6273 | 2989 | 430 |
| | Macrophage | 21162 | 7462 | 426 |
| | Endothelial | 2810 | 1069 | 496 |
